# Strong genetic differentiation but limited niche partitioning in a sympatric species pair separated by an allochronic reproductive barrier

**DOI:** 10.1101/2023.08.30.555036

**Authors:** Mitchell Irvine, Zachary Stewart, Nagalingam Kumaran, Chapa G. Manawaduge, James Ryan, Solomon Balagawi, Brendan Missenden, Melissa Starkie, Anthony R. Clarke, David Hurwood, Peter Prentis

## Abstract

The combination of differential adaptation to ecological niches and the development of reproductive barriers are considered helpful for maintaining co-existing species. In the absence of one of these elements, species boundaries are expected to breakdown. The tephritid fruit fly species pair, *Bactrocera tryoni* and *B. neohumeralis*, have significant overlap in geographic range and host use, with time of day of male mating the only known difference in their mating systems. Using a combination of ecological (seasonal abundance, host use and habitat use) and genomic data, we tested the differing roles of competition and assortative mating on the maintenance of the species boundaries in this species pair. Genome-wide SNP analyses found strong genetic differentiation between the species with no evidence for hybridization in the field. Most outlier SNPs were restricted to narrow regions towards the centromeres and telomeres of chromosomes and high nucleotide diversity rates were observed throughout the chromosomes of both species. Enrichment of annotation terms indicated an overabundance of genes with the ‘abnormal neuroanatomy’ term. Terms of interest associated with sleep and circadian rhythm, potentially important to the allochronic reproductive barrier, were non-enriched. Ecological data found limited evidence for ecological differentiation between the two species based on significant positive correlations between species numbers trapped at different times of the year, trapped in different habitats within a region, or when reared from fruit.

Our study highlights the significance of assortative mating over ecological differentiation for sympatric maintenance of the *B. tryoni*/*B. neohumeralis* sibling pair.

## Introduction

Understanding the ecological and genetic basis of speciation and species maintenance is a fundamental element of evolutionary biology. In geographically isolated populations, an absence of gene flow between incipient species is expected to allow for the accumulation of genetic divergence via both selection and drift (Feder et al., 2013). When two populations experience gene flow (either through primary or secondary contact), reproductive isolation is considered necessary to limit its homogenizing effect (Coyne & Orr, 2004).

Central to our understanding of speciation in the presence of gene flow are the opposing roles of natural selection and recombination on the formation or dissolution of species boundaries (Felsenstein, 1981). Natural selection promotes the development of reproductive isolation through the formation of locally adapted gene complexes, while free recombination acts to dissolve them through breaking down the associations of such complexes (Felsenstein, 1981; Hey, 2001; Schluter & Rieseberg, 2022); a reduction in recombination rates is considered helpful for species boundaries to persist and strengthen over time (Butlin, 2005). Traits associated with reproductive isolation, such as those involved in assortative mating, are considered effective mechanisms for constraining gene flow and are essential for sympatric species divergence when coupled with the development of traits associated with local adaptation (Ortiz-Barrientos et al., 2016; Schluter & Rieseberg, 2022).

On an ecological scale, the persistence of two species in sympatry depends on two criteria; their ability to overcome the effects of hybridization through the development of reproductive isolation, and the development of niche partitioning to avoid inter-species competition asymmetries (Coyne & Orr, 2004; Felsenstein, 1981; Weber & Strauss, 2016). Without meeting these criteria, theory suggests that species will likely dissolve through hybridization or cease to exist in sympatry through competitive displacement (Germain et al., 2021; Weber & Strauss, 2016). Although the coexistence of ecologically equivalent species may occur long term where sexual selection exists, provided the local carrying capacity is sufficient enough to maintain both populations (M’Gonigle et al., 2012).

The Australian frugivorous tephritids (Diptera: Tephritidae), *Bactrocera tryoni* (Froggatt) and *Bactrocera neohumeralis* (Hardy), are a fruit fly sibling pair, studied for their speciation histories and genetic stability for over 60 years (Birch, 1961; Gilchrist et al., 2005; Gilchrist & Ling, 2006; Gilchrist et al., 2014; Lewontin & Birch, 1966; Meats et al., 2003; Pike, 2004; Raphael et al., 2019; Smith, 1979; Yeap et al., 2020). Genetically, *B. tryoni* and *B. neohumeralis* are very similar (Gilchrist et al., 2014; Morrow et al., 2000; Wang et al., 2003), with shared polymorphisms in coding and non-coding regions indicating continued genetic exchange or recent separation (Gilchrist et al., 2005; Gilchrist & Ling, 2006; Wang et al., 2003). The species show only minor variation in their morphology; the humeral calli, located on the ‘shoulders’ of the fly, are bright yellow in *B. tryoni* and brown in *B. neohumeralis* (Drew, 1989). While intermediate colour variants are well documented, genetic analysis using microsatellites suggest that intermediates represent intraspecific phenotypic variation, rather than hybrid individuals (Gilchrist & Ling, 2006; Pike, 2004; Wolda, 1967). While *B. tryoni* is typically more abundant than *B. neohumeralis*, the geographic ranges of the two species extensively overlap along the east coast of Australia, with the endemic range of *B. neohumeralis* entirely encompassed by the range of *B. tryoni* except for a few islands in the Torres Strait, north of Queensland (QLD) (Fig. 1A,B) (Osborne et al., 1997). Both *B. tryoni* and *B. neohumeralis* are highly polyphagous, using fruit from multiple plant families for breeding, with host utilization data [presented as ‘simple’ lists of host fruit that either species have been reared from at least once with little or no further information] showing significant overlap between the pair (Fig. 1C) (Hancock et al., 2000; Leblanc et al., 2012; QDPC, 2021).

**Figure 1.**
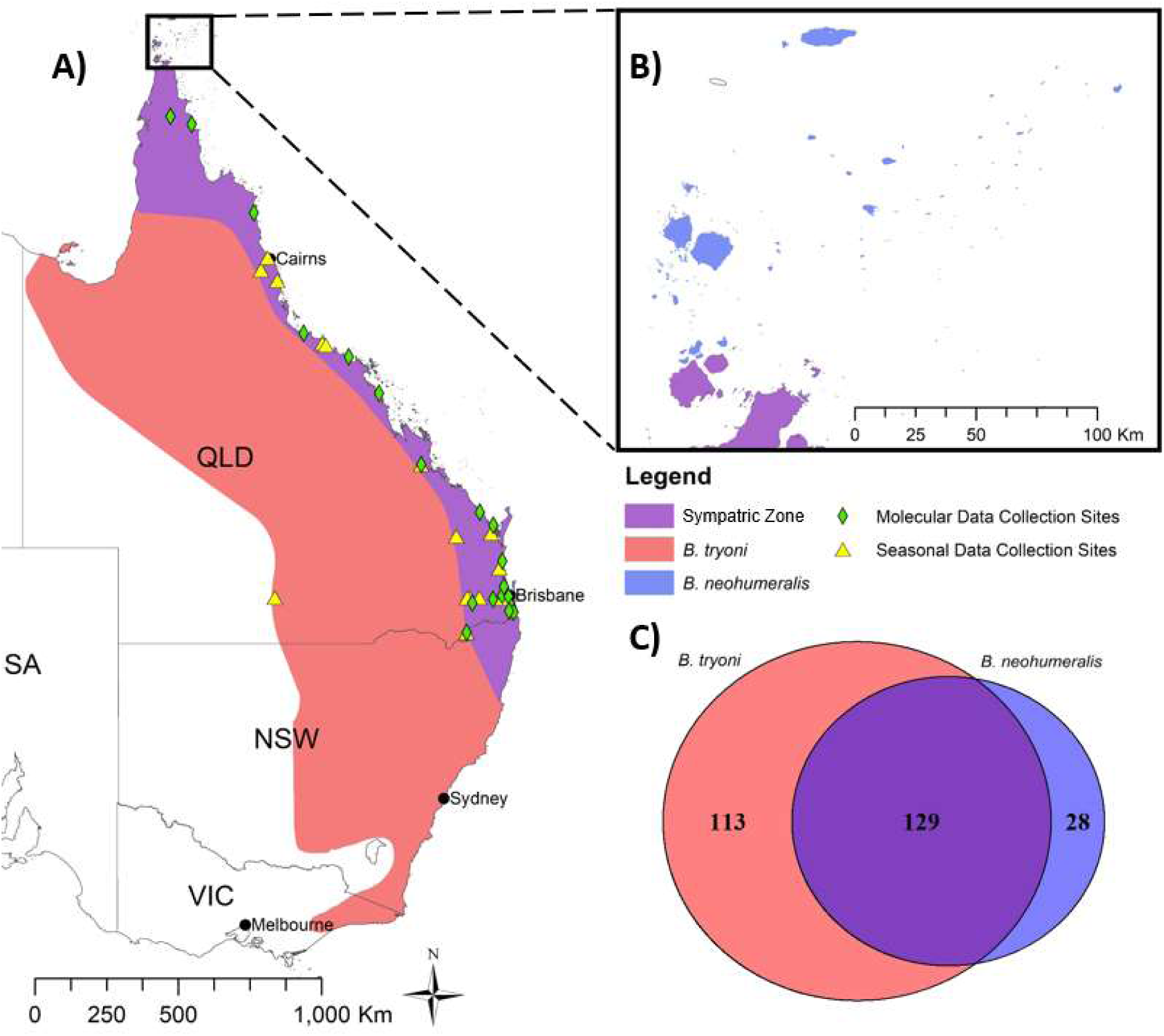
A) Distribution of *Bactrocera tryoni* and *B. neohumeralis* along the east coast of Australia. B) Distribution of both species in the Cape York Peninsula and Torres Strait Islands. C) Host usage overlap between species. Numbers in circles represent the known number of host species utilized. Range and host data gathered from Hancock et al. (2000); Leblanc et al. (2012); QDPC (2021). Note: Recent expansion during the past 50 years of *B. tryoni* into central and western Victoria is not shown because it is still under regulatory control in many places and is not associated with the species’ evolutionary history.

So closely related are *B. tryoni* and *B. neohumeralis* that only one known trait defines them as separate biological species: their time of mating (Birch & Vogt, 1970; Lewontin & Birch, 1966; Vogt, 1977). *Bactrocera neohumeralis* mates during the middle of the day at high light intensity over a mating window of three to seven hours, while *B. tryoni* initiates mating during dusk at low light intensity in a mating window of approximately one hour (Ekanayake et al., 2017). This difference in mating time is the only known mechanism maintaining reproductive isolation between the species (Birch & Vogt, 1970; Lewontin & Birch, 1966; Meats et al., 2003; Raphael et al., 2019; Smith, 1979), with other components of their mating system not known to be different (Bellas & Fletcher, 1979; Ekanayake et al., 2017; Mankin et al., 2008), including an absence of post-zygotic barriers (Yeap et al., 2020). Female *B. neohumeralis* will, like *B. tryoni*, also mate at dusk, at which time *B. tryoni* males do not discriminate between females of either species (Yeap et al., 2020). This caused Yeap et al. (2020) to conclude “*the mating time difference may be a weaker reproductive isolating barrier than once assumed*”.

If mating time is a weak reproductive barrier between the two species, then other factors must be playing a role in maintaining separate species; for example, genomic incompatibility (Dobzhansky, 1936; Turelli et al., 2001). Additionally, for herbivorous insects such as fruit flies, species divergence is largely considered to be driven by host switching and host specialization that reduces the effects of competition and natural enemies (Gavrilets & Losos, 2009; Janz & Nylin, 2008; Jeffries & Lawton, 1984; Schluter, 2000; Yoder et al., 2010) and so ecological factors also need to be considered when investigating the maintenance of species boundaries. Conversely the species may not be evolutionarily stable, in which case evidence of gene flow should be expected from sympatric field collections of the two taxa.

In this study, we conducted a comprehensive investigation of how the lineages of *B. tryoni* and *B. neohumeralis* species have diverged and been maintained in apparent sympatry through dense geographic sampling of nuclear genomes, and structured field sampling of temporal population abundance and habitat-and host-usage. We assessed the extent of ongoing gene flow and genetic differentiation between the species and identified putative candidate genes associated with reproductive isolation. Using the field data, we conducted correlation analyses on seasonal abundance, host use and habitat utilization for evidence of competitive displacement between the two species. Shifts in any or all of these three ecological attributes have been previously associated with competition between fruit fly species (Clarke & Measham, 2022).

## Methods

### Molecular data sample collection

Individuals of *B. tryoni* and *B. neohumeralis* were collected in December 2020 and December 2021, using cue lure baited Steiner (Steiner, 1957), Lynfield (Cowley et al., 1990) or Paton traps (Plant Health Australia, 2023). These traps were deployed by Queensland Department of Primary Industries (QDPI) as part of ongoing Biosecurity Queensland (BQ) surveillance. A subsample of the trapping sites was selected from the existing network to represent the geographic range of both species (Fig. 1A). Up to 10 individuals of each species were collected per site with a total of 81 *B. tryoni* and 105 *B. neohumeralis* individuals collected from 19 sites. Detailed trapping records are included in Table S1. Identity of the samples were confirmed following the descriptions in fruit fly identification handbooks (Drew, 1989; Plant Health Australia, 2018), and samples were stored at −20°C in 100% ethanol.

### Extraction and sequencing

DNA was extracted using the QIAGEN DNeasy® Blood & Tissue Kit following the manufacturer’s protocol. Samples were screened for quality and quantity, using both gel electrophoresis and Qubit assay, and then sent to Diversity Arrays Technology Pty Ltd, Canberra (DArT P/L), for DArTseq high-density genotyping (Kilian et al., 2012). A PstI/SphI, restriction enzyme combination was used by DArT P/L, and the fragments (up to 90bp) were sequenced on an Illumina Hiseq2500 as single end reads.

### Data sourcing

All relevant analyses made use of the *B. tryoni* genome assembly (Genbank accession = GCF_016617805.1).

### Read alignment and SNP calling

Reads were demultiplexed using process_radtags in Stacks v2.60 (Rochette et al., 2019). Read mapping used BWA-MEM (Li, 2013) to produce a SAM alignment file with appropriate read group specification for each sample. Subsequently, SAMtools (Li et al., 2009) produced sorted and indexed BAM alignment files. Freebayes v1.3.6 (Garrison & Marth, 2012) was used in an iterative process to predict single nucleotide polymorphisms (SNPs) in all samples. Firstly, each sample had SNP calling performed individually using freebayes; output Variant Call Format (VCF) files were left-aligned using BCFtools (Danecek et al., 2021) and block substitutions were decomposed using vt (Tan et al., 2015). Resulting VCF files were merged using BCFtools, and this VCF file was then used as input for a second round of SNP calling using freebayes (*-@ file.vcf --only-use-input-alleles*) to ensure all samples were genotyped at the same locations. The variant calls were filtered using a method based upon James et al. (2021) wherein SNPs were removed if they had missing data in > 50% of the population, had a quality score < 30, a depth < 3 read alignments, or a minor allele count < 1; we also filtered out SNPs with a minor allele frequency < 5%.

### Population structure and genetic differentiation

To investigate whether admixture was occurring between the two species, we used fastSTRUCTURE v1.0 (Raj et al., 2014). A simple prior of K=1-10 was used, after which the chooseK.py utility program provided with fastSTRUCTURE was used to assess the optimal number of genetic clusters to explain the structure among species and populations. Plots were generated using pophelper v2.3.1 (Francis, 2017) in R.

Additionally, we ran a Principal Component Analysis (PCA) to provide further visualization of the variations within and between the two species, using the SNPRelate v1.24.0 (Zheng et al., 2012) package in R. The linkage disequilibrium-based SNP pruning function provided with this software was run with default parameters to mitigate any bias associated with linked SNP clusters upon the downstream PCA. We similarly performed this SNP pruning prior to calculation of IBS proportions for samples using SNPRelate.

### Relatedness prediction

Relatedness analyses were performed in R using the dartRverse package (Gruber et al., 2018; Mijangos et al., 2022). Variants were filtered to remove those with a callrate lower than 95%, those deviating from Hardy-Weinberg Equilibrium to a significance threshold of 0.05, and those with linkage disequilibrium values above 0.2. The remaining variants were used as input for the EMIBD9 (Wang, 2022) and rrBLUP (Endelman, 2011) packages to estimate the relatedness of samples in a pairwise manner.

### Population genetics statistics

Mean FST (θ) values (Weir & Cockerham, 1984) were calculated both within and between the two species using VCFtools (Danecek et al., 2011). Within species comparisons were conducted for each species separately and samples from each collection site were considered separate populations. For between species comparison, the samples were grouped by species and entered as two separate populations.

Additional population statistics were calculated to estimate levels of relative genetic differentiation between species (F_ST_), absolute genetic divergence between species (dXY), and nucleotide diversity within species (π). These statistics were calculated within non-overlapping windows of 50 Kbp using pixy (Korunes & Samuk, 2021). As pixy explicitly requires invariant sites to be called, we recalled variant and invariant sites using a pipeline involving bcftools mpileup (-p -A -d 500) followed by bcftools call (-m), with resulting predictions used as input. These statistics were mapped along each chromosome to identify patterns of differentiation within the chromosomal landscape.

### Recombination rate prediction

The recombination rate along each chromosome was approximated using a recurrent neural network as implemented by ReLERNN (Adrion et al., 2020). The variants for each species were filtered to retain only biallelic SNPs, after which each species’ data was processed using ReLERNN_SIMULATE to generate training and validation data (with default settings), ReLERNN_TRAIN to train the neural network (with --nEpochs 100 and --nValSteps 20), ReLERNN_PREDICT to predict per-base recombination rate, and ReLERNN_BSCORRECT to perform parametric bootstrapping and estimate 95% confidence intervals.

### Population structure along chromosomes

Population structure was assessed within local regions of each chromosome as implemented by the LOSTRUCT method (Li & Ralph, 2018). The run_lostruct.R wrapper script (available within the program’s GitHub repository at https://github.com/petrelharp/local_pca) was configured to assess local structure using PCA within non-overlapping windows of 50 SNPs. For each window, the first principal component was extracted for each sample and the spread of these values were visualized by their median and the 25% and 75% quantiles to identify regions where population structure was more apparent.

### Outlier SNPs and gene candidates

Variant predictions were converted from VCF to GESTE format using PGDSpider v2.1.1.5 (Lischer & Excoffier, 2012), then outlier SNPs were predicted using Bayescan v2.1 (Foll & Gaggiotti, 2008) with default parameters. Genes containing outlier SNPs within their gene model (exons and/or introns), as well as genes that were the nearest to an intergenic outlier SNP were identified using a custom script.

Candidate genes were queried against the *Drosophila melanogaster* gene models (Larkin et al., 2021) (FlyBase release FB2023_01) using MMseqs2, and the FlyBase automated gene summaries of the best hit were parsed for functional and phenotypic annotations which we attributed to sequences if a minimum E-value of 1e^-10^ was achieved.

### Seasonal abundance

A historical dataset (May, 1961) was used for the comparison of temporal abundance of *B. neohumeralis* with *B. tryoni*. This dataset has been previously used to report *B. tryoni* [but not *B. neohumeralis*] seasonal phenology (Muthuthantri et al., 2010), and so the phenology patterns *per se,* are not explored here. Rather, we focus on evidence for temporal niche segregation in *B. neohumeralis* and *B. tryoni* through paired t-tests and Pearson correlation analysis using the R package ggpubr (Kassambara, 2023). In Africa, a shift in seasonal abundance to minimize an overlap in host usage (i.e. temporal niche displacement) has been reported as the outcome of competition between the native mango fruit fly (*Ceratitis cosyra* (Walker)) and invasive Oriental fruit fly (*Bactrocera dorsalis* (Hendel))(Vayssières et al., 2015). Therefore, we predict that if there is competition-driven temporal niche displacement between the two species, then there should be no, or even negative correlation between their seasonal abundances. We interpret significant positive correlation between their populations as a lack of evidence of temporal niche displacement.

Detailed methodology on the fly trapping dataset is described elsewhere (May, 1961; Muthuthantri et al., 2010). In summary, 15 trapping sites across Queensland, Australia, were used, ranging from temperate Stanthorpe in the south to tropical Cairns, ∼1500 km to the north (see Fig.1A and Table S5). Fly trapping was carried out for up to six years per location using a fruit-based liquid lure trap (May & Caldwell, 1944). Ten traps were maintained at each location, with traps cleared weekly. The data in May (1961) vary between sites in how they are presented, and modifications were needed in order to directly compare *B. tryoni* and *B. neohumeralis* populations. Six sites were excluded as seasonal data were not available (see Table S5 for details). The t-tests and correlation analyses between *B. tryoni* and *B. neohumeralis* populations were conducted on a site-by-site basis (i.e., data were never collated across sites), so the different manipulations performed on the site-specific data sets did not affect results.

Given the variability in how data from different locations are presented in the historical data, a dataset systematically collected as part of a recent landscape genetics study was also used to assess seasonal abundance on a fine scale. Detailed sampling techniques are presented in Ryan et al. (In Review) with abundance data being used for the seasonal abundance correlation analysis and the landscape ecology analysis below. In brief, samples were collected from 25 locations in the Wide Bay-Burnett Region of Queensland (see Fig. 2) in April 2021, August 2021, October 2021, December 2021, February 2022 and April 2022. The t-tests and correlation analyses between *B. tryoni* and *B. neohumeralis* populations was conducted for each bi-monthly collection.

**Figure 2:**
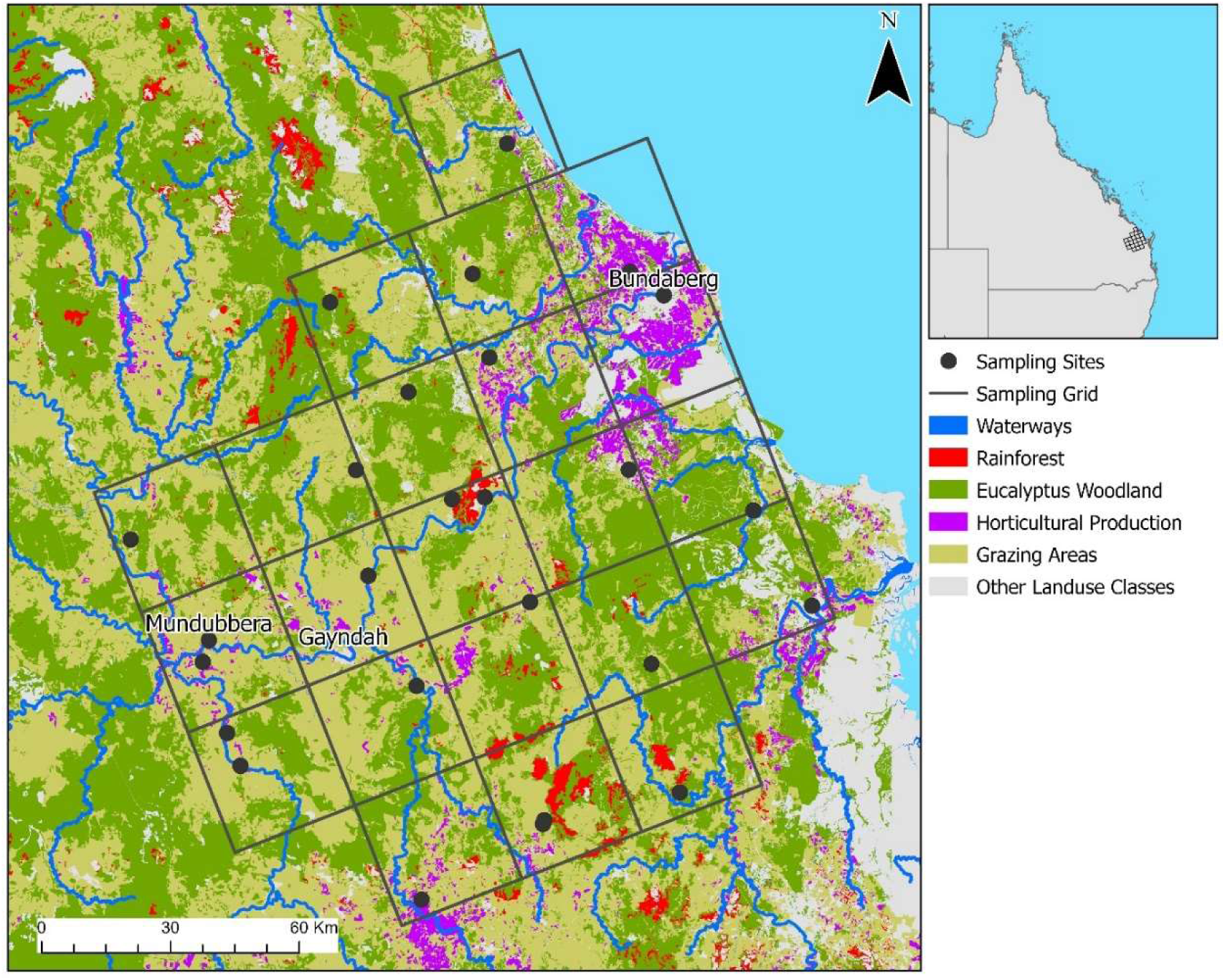
Collection sites throughout the Wide Bay-Burnett region of South East Queensland for the fine scale seasonal abundance analysis. Taken from Ryan et al. (In Review).

### Habitat use

This study compared abundance of *B. tryoni* and *B. neohumeralis* across different human-defined habitat types to determine habitat use patterns. Six habitat types, grassland, suburbia, horticultural farming, mixed sugarcane and other crops (mixed farming hereafter), dry forest, and wet forest were chosen for the study in the Bundaberg region of Queensland (∼350 km north of Brisbane). Specific descriptions of each habitat type are outlined in Table S7. For each habitat type there were nine replicate fruit fly traps. Trapping commenced in September 2010 (spring) and was completed in March 2011 using cuelure baited modified Steiner traps (Drew et al., 1982). The trapping period covers the period of peak yearly activity for *B. tryoni* (Clarke et al., 2022). The trapping program involved setting up the traps on a monthly basis at each trap site, removing the traps at each site three days after setting up and replacing the traps with fresh wicks and attractants in the following month. Flies collected over the three days in each month were counted and identified to species level using a fruit fly identification guide (Drew, 1989). Paired t-tests and Pearson correlation analyses on the total trapping dataset for a habitat type were performed to compare the relationship in abundance of the two different species at that habitat type. Examples of spatial displacement of competing fruit flies are presented in Duyck et al. (2004).

### Landscape ecology

Linear mixed modelling (LMM) was used to investigate patterns in *B. neohumeralis* abundance across the study area (See Fig. 2) to complement a study conducted by Ryan et al. (In Review) which investigated the environmental predictors of *B. tryoni* abundance. Additionally, the Empirical Bayesian Kriging (Geostastical Analyst Tools) tool was implemented in ArcGIS Pro v3.4 to interpolate *B. neohumeralis* abundance across the study area to visualise spatially explicit patterns of abundance.

The *B. neohumeralis* abundance data collected throughout the Wide Bay-Burnett region described in the fine scale seasonal abundance analysis above was used. The LMM analysis incorporated the same landscape predictor variables as was used in the LMM analysis in Ryan et al. (In Review), briefly, % coverage of five landscape categories (grazing area, residential area, *Eucalyptus* woodland, rainforest and agricultural production) within a 1km radius of each sampling site were investigated. Additionally, other variables investigated included latitude, longitude, and distance from each sampling site to the nearest perennial water source.

Based on the LMM analysis methods outlined in Ryan et al. (In Review), the R package lme4 version 1.1-30 (Bates et al., 2015) was used to create our LMMs and ‘dredge()’ function implemented in the R package MuMIn version 1.47.1 (Barton, 2022) to assess all combinations of predictor variables, using sampling time as a random variable. Following dredging, LMMs with an ΔAICc < 2 were used to perform model averaging and determine for each landscape predictor; linear coefficient, standard error, statistical significance and upper and lower 95% confidence intervals.

### Host fruit use

Fruits were collected from the QDAF Maroochy Research Facility, Nambour, Queensland (∼100 km north of Brisbane) from separate orchard blocks that consisted of either mixed tropical fruit varieties, stone fruit or mangoes, as well as some isolated trees. Small numbers of ripe or mature green fruit were sampled on a weekly or fortnightly basis between December 2012 and October 2015. Fruit were selected from a number of trees within the orchard block and picked from various heights and aspects to obtain a random sample. Fruits collected were: white sapote (*Casimiroa edulis*), mulberry (*Morus nigra*), peach (*Prunus persica*), plum (*Prunus domestica*), carambola (*Averrhoa carambola*), nectarine (*Prunus persica*), guava (*Psidium guajava*), sapodilla (*Manilkara zapota*), *Syzigium* spp., grumichama (*Eugenia brasiliensis*), hog plum (*Spondias mombin*), jabotica (*Plinia cauliflora*), white mulberry (*Morus alba*), avocado (*Persea americana*), black sapote (*Diospyros nigra*), longan (*Dimocarpus longan*), mango (*Mangifera indica*) and cashew (*Anacardium occidentale*).

Fruit were placed in paper bags and transported to the QDAF Brisbane laboratories. Fruit were counted and weighed and then placed on gauzed plastic containers over vermiculite, in plastic boxes with gauzed lids to allow ventilation. Boxes were held in a Controlled Environment Room (26°C and 70%RH) to allow insects to develop through to the pupal stage. The vermiculite was sieved weekly until all insects had exited fruit and pupated, and fruit were inspected before being discarded. Fruit fly pupae that were reared out of fruit samples were placed into small plastic boxes with gauzed lids containing vermiculite. Once adult fruit flies had emerged from pupation, they were identified to species level using morphological taxonomic characters. The abundance of the two fly species from a host was compared using a paired t-test and Pearson correlation analysis. Changes in fruit fly host usage patterns due to competition are well documented in the fruit fly literature (Charlery de la Masselière et al., 2017; Hassani et al., 2022).

## Results

### Molecular Data

#### Read processing, alignment and SNP calling

After demultiplexing samples, between 2,596,430 and 5,228,647 high-quality sequence reads were obtained for each sample. Alignment to the *B*. *tryoni* genome assembly using BWA-MEM indicated that most samples had high mapping rates (median alignment percentage = 74.98%), with a small number of samples having low sequence alignment (minimum alignment percentage = 0.38%); see Table S2 and Table S3 for full details of read counts and alignment percentages. 28,334 SNPs were identified using Freebayes from the two species that met all predetermined threshold criteria for inclusion.

#### Populations cluster by species and not geography

We found that populations clustered based on species and not geography in all analyses. PCA (Fig. 3) and fastSTRUCTURE (Fig. 4) both resolved species-specific groups, demonstrating clear genetic delineation of the two species and an absence of intermediate genotypes between the species. Several *B. neohumeralis* samples (genotype codes 526, 528, 529, 530, and 531 from the NS location) appear to be outliers based on PCA (See Fig. S1), but one sample from the same location (527) grouped with the majority of *B. neohumeralis* samples in ordination space. Additionally, the chooseK.py function of fastSTRUCTURE predicted three population clusters, with the outlier samples from the PCA representing the third cluster (See Fig. S2).

**Figure 3.**
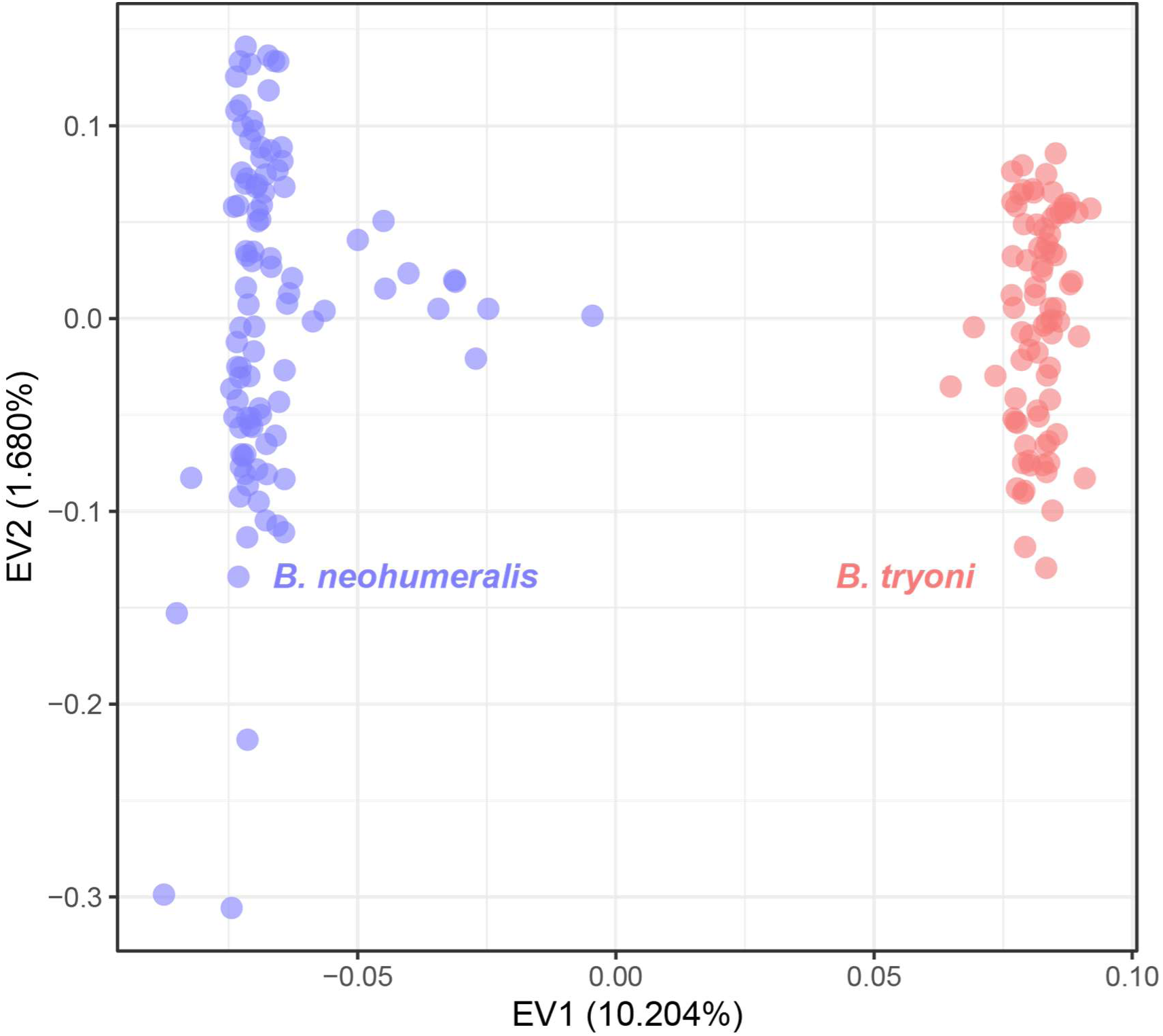
Principal component analysis (PCA) of single nucleotide polymorphisms (SNPs) *Bactrocera neohumeralis* and *B. tryoni*, with the values from the first two eigenvectors plotted for each individual. Eigenvectors one (horizontal) and two (vertical) explain 10.204% and 1.68% of the variance in the data, respectively.

**Figure 4.**
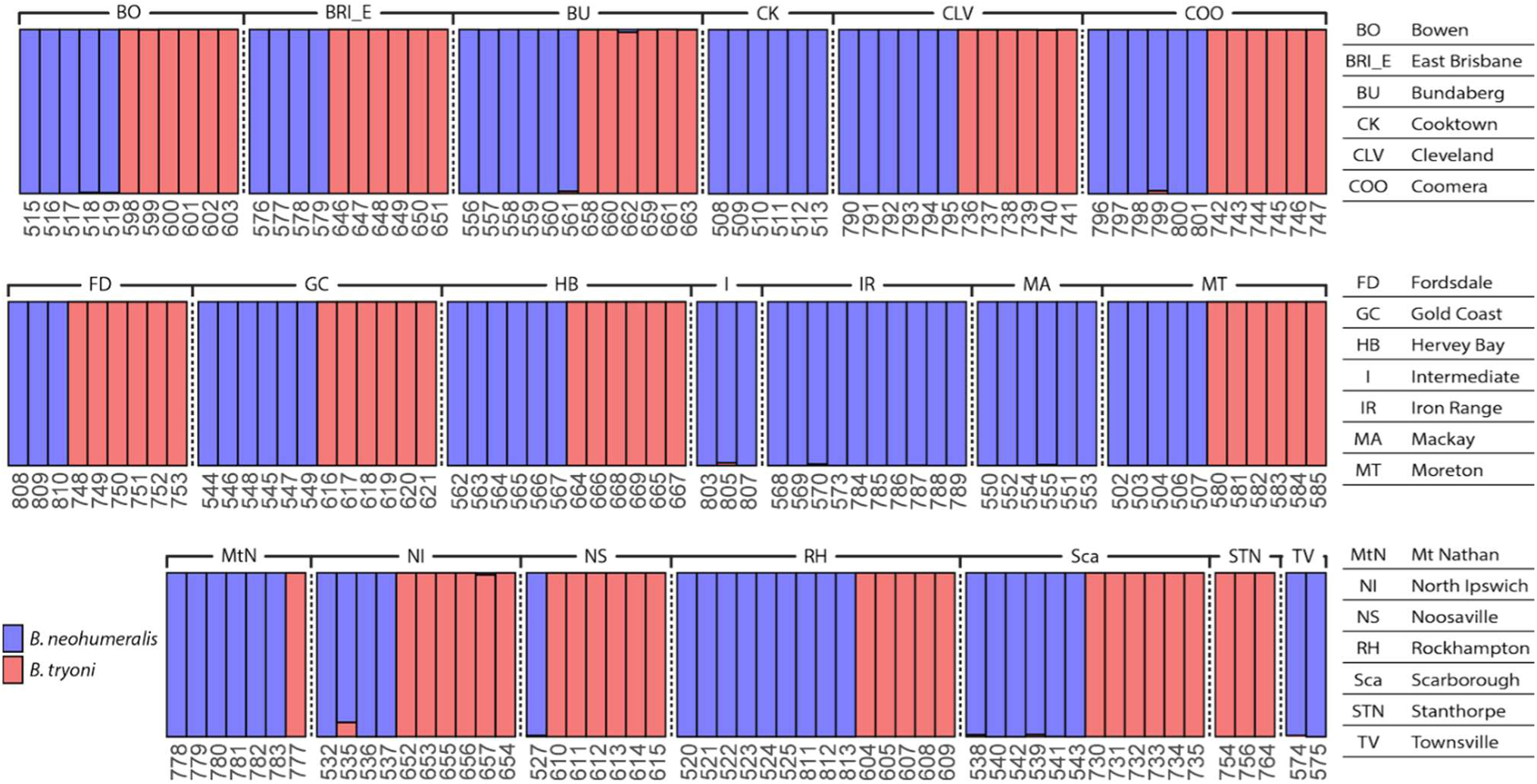
fastSTRUCTURE results at cluster size (K) equals two for single nucleotide polymorphisms (SNPs) for *Bactrocera neohumeralis* and *B. tryoni* grouped by collection site. Red and blue colouration of the bars indicates the inferred population admixture proportions as a percentage (numbers not shown). Three-digit numbers along the bottoms of each row of bars are genotype ID codes. The intermediate grouping (I) includes samples with an intermediate phenotype. All three intermediate samples (genotypes 803, 805 and 807) presented as *B. neohumeralis* genotypes.

These outlier samples had poor genotype calling rates (min = 27.9%, max = 32.3%) compared to sample 527 (91.3%) and most other samples (median = 89.1%). As such, we believe these outliers to result from sequencing issues and were excluded from further analysis.

#### Relatedness of individuals

After filtering, 3710 variants remained for use in the relatedness analyses. The EMIBD9 estimates of relatedness (i.e., Identity by Descent) for the sampled individuals are presented in Figure 5. Notably, each species is largely segregated into distinct groups showing increased relatedness values within species groups compared to between species groups where values approached zero. Three samples (genotype codes 662, 787, and 788) appear as an outgroup relative to the other samples and show elevated levels of relatedness both within the same species and between species. While the levels of relatedness are elevated in these samples, they still present higher levels of relatedness within species groups compared to between species groups. These samples do not appear as anomalies in any other analysis suggesting a lack of data after filtering preventing the EMIBD9 analysis from discerning these samples resulting in elevated levels of relatedness.

**Figure 5.**
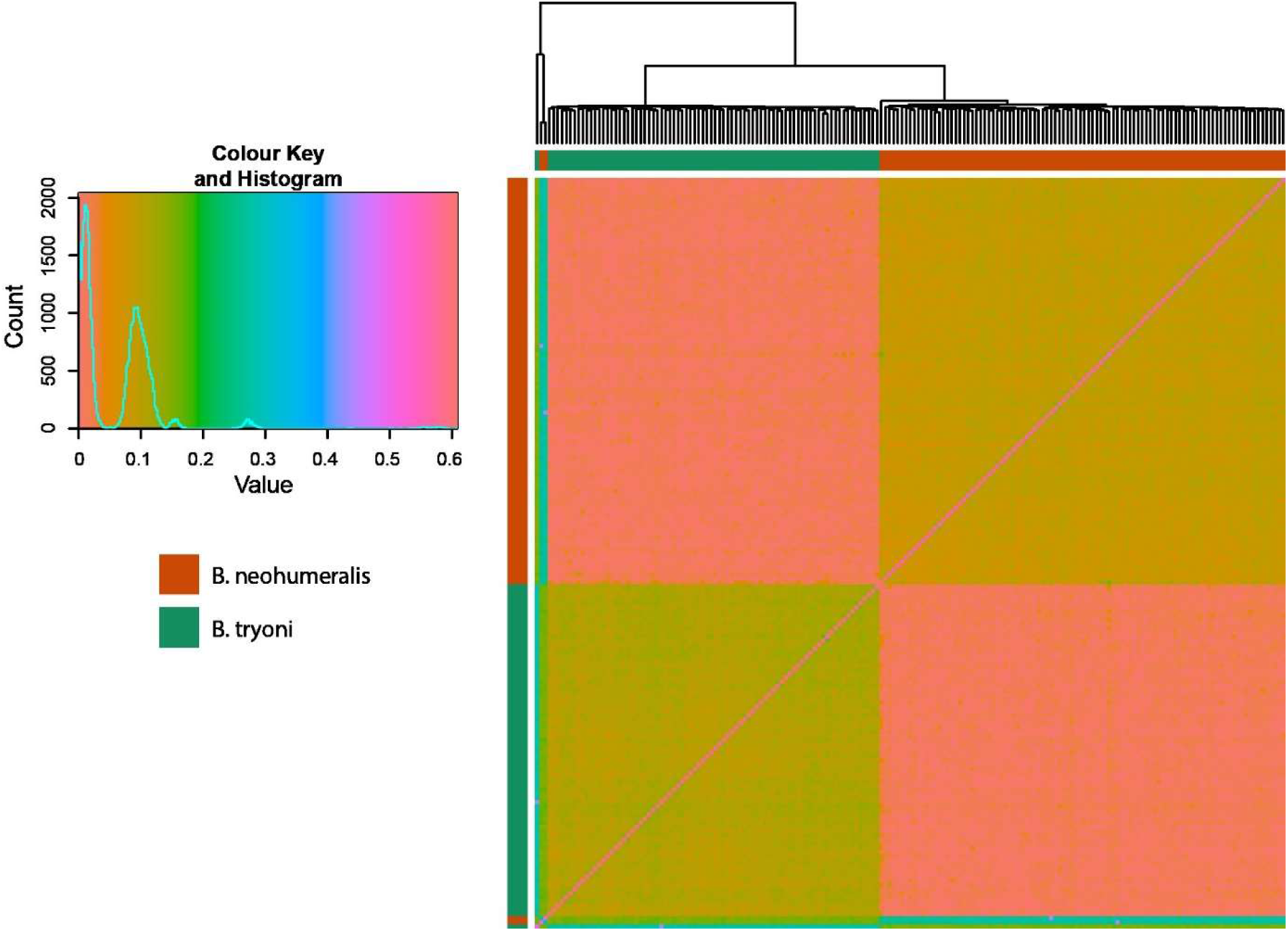
Heatmap generated using the results of EMIBD9 assessing the relatedness of sampled individuals via their Identity by Descent (IBD) coefficients. The colour key indicates the coefficient value with a histogram plotted on top of the colour key to showcase the amount of pairwise sample comparisons obtaining that value. The heatmap indicates the IBD value for each pairwise sample comparison, with the sample’s species indicated along the left and top of the plot. Lower values (e.g., soft red colour) indicate low to no relatedness as seen when comparing most *Bactrocera neohumeralis* samples against most *B. tryoni* samples. An increased value (e.g., amber colour) indicates some degree of relatedness, and is seen when comparing most species amongst themselves.

The rrBLUP estimates of relatedness are presented in Figure 6. Notably, genotype codes 662, 787, and 788 do not appear to be outliers in this analysis unlike what was seen in the results of EMIBD9; all species only show relatedness to other members of their species group with negative values obtained when compared to the opposite species suggesting non-random mating between the species groups.

**Figure 6.**
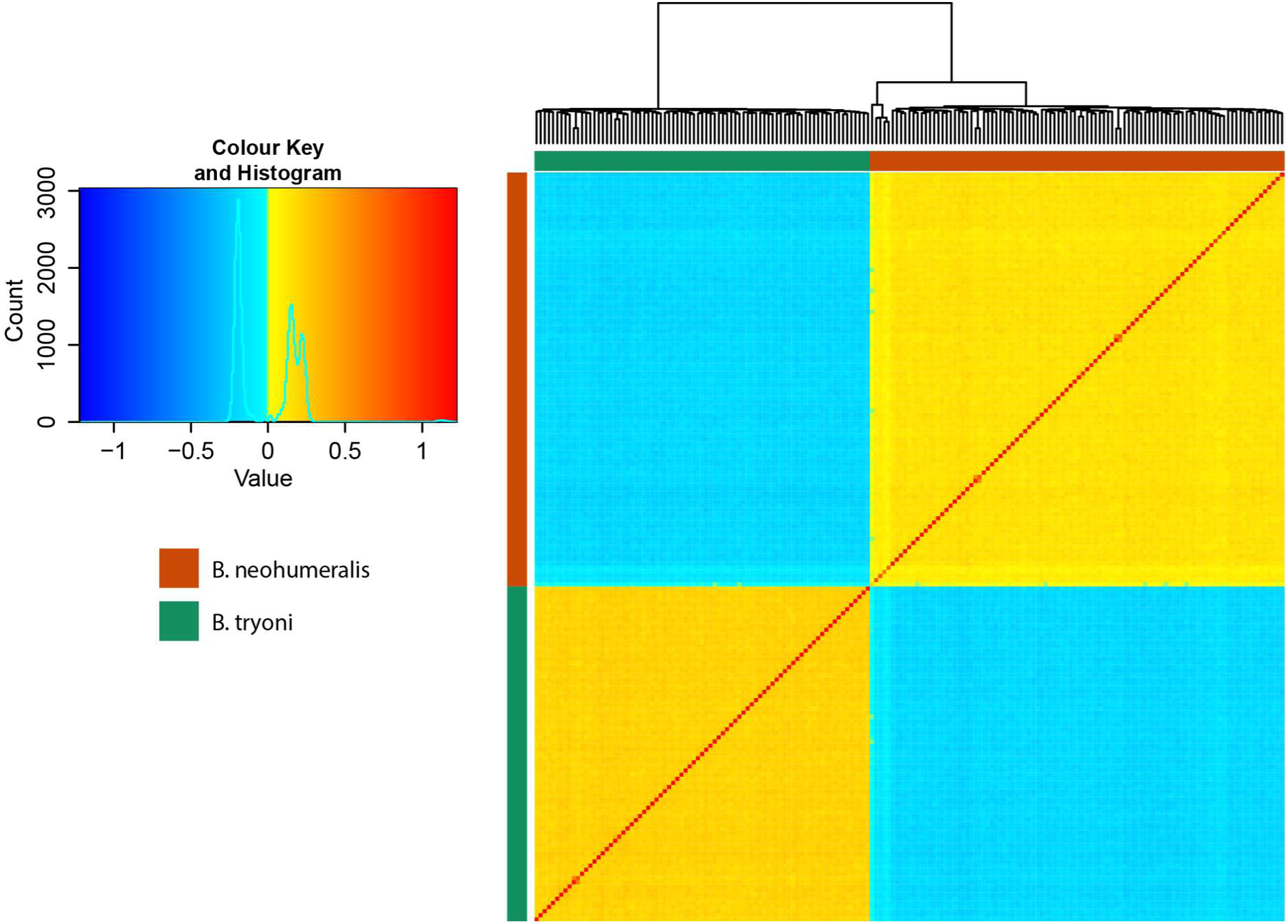
Plot generated using the results of rrBLUP assessing the estimated relatedness of sampled individuals via an additive relationship matrix. The colour key indicates the probability of identity by descent with a histogram plotted on top of the colour key to showcase the amount of pairwise sample comparisons obtaining that value. A heatmap indicates the estimated relatedness value for each pairwise sample comparison, with the sample’s species indicated along the left and top of the plot. Negative values (e.g., blue colour) indicate low to no relatedness as seen when comparing *Bactrocera neohumeralis* samples against *B. tryoni* samples. A positive value (e.g., yellow colour) indicates some degree of relatedness and is seen when comparing individuals within a species.

#### Population genetics statistics, recombination rates and structure along chromosomes

Patterns of differentiation among populations (F_ST_) were also lower between populations within a species (*B. neohumeralis* = 0.0025796 when ignoring outlier samples, *B. tryoni* = 0.0036269) and two orders of magnitude higher between species (0.10843). Using outlier detection in BayeScan, 357 of the 28,334 SNPs (1.26%) displayed a significantly elevated level of differentiation between the species. The distribution of outlier SNPs along autosomal chromosomes (See Fig. 7 to Fig. 11) was largely restricted to ∼ 16 narrow genomic regions. Elevated windowed F_ST_ and d_XY_ values calculated using Pixy were also clustered in similar regions to those observed in the outlier SNPs. These narrow genomic regions with outlier loci were not uniformly distributed across the five autosomal chromosomes with 10 occurring on chromosome ends and two near the centre of chromosomes. A relatively small number of outlier SNPs were mapped to ‘unplaced contigs’ (See Fig. S3). Identity-By-State (IBS) analysis also provided insight into the genetic homogeneity of the two species, with *B. tryoni* having a mean value of 0.797 and *B. neohumeralis* having a mean value of 0.790 (species with a value closer to 1 are more similar) suggesting that both species are approximately equally homogenous.

**Figure 7.**
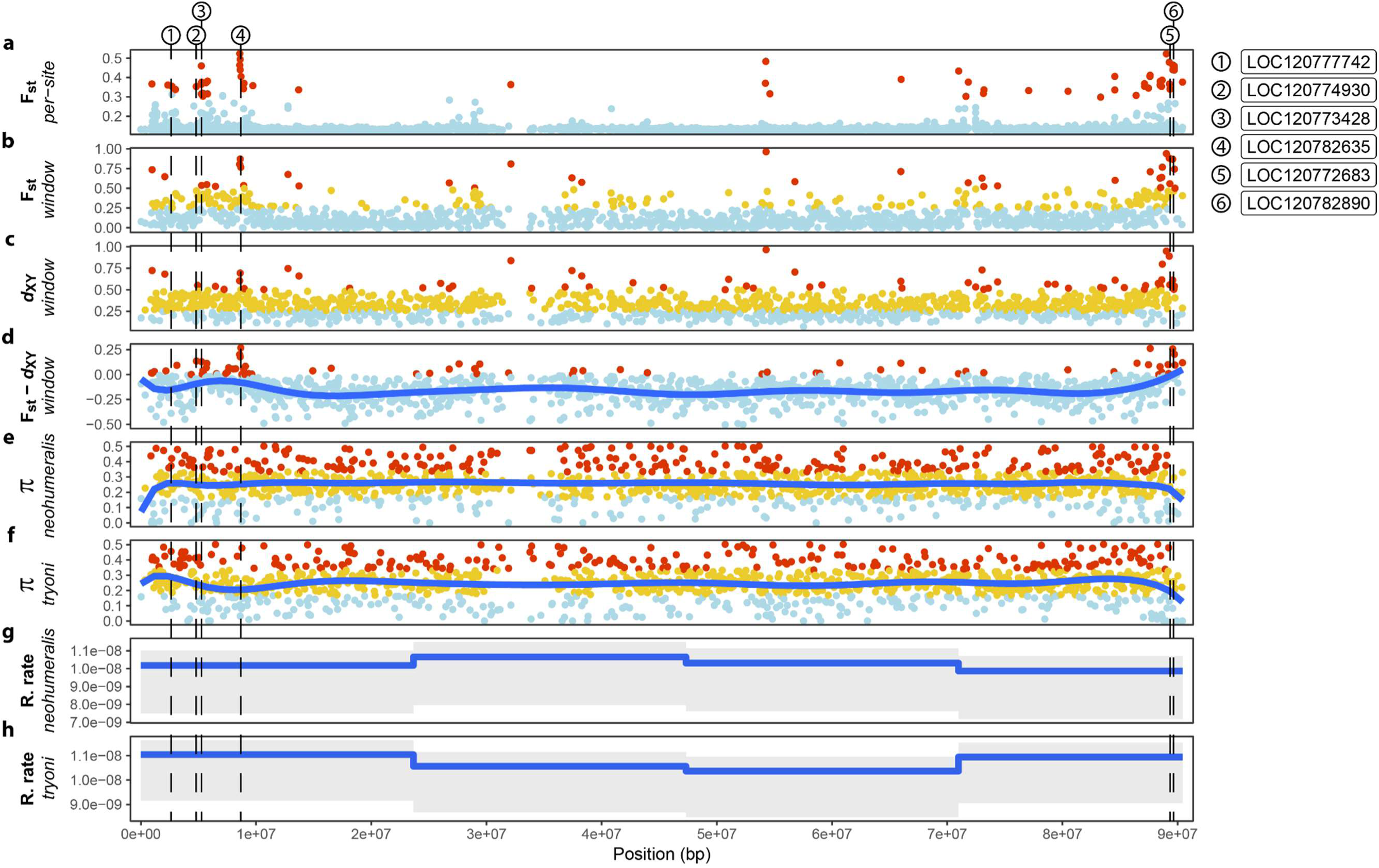
Population genetics statistics calculated for chromosome 1 labelled as NC_052499.1. Where indicated on the Y-axis that a statistic is windowed, values have been obtained through averaging over window lengths of 50 Kbp. Each statistic is presented along the length of the chromosome where the position labelled as 1e+07 equates to 10 Mbp. Candidate genes potentially involved in the assortative mating of the two species are indicated as gapped vertical lines denoted by a number which relates to the locus identifiers shown in the upper right corner of the figure (See Table S4 for candidate gene details). For subfigures b and c, colours are depicted according to values being in the range of: 0 to 0.25 is light blue; >0.25 to 0.5 is yellow; >0.5 to 1 is red. For subfigures e and f, colours are depicted according to values being in the range of: 0 to 0.2 is light blue; >0.2 to 0.35 is yellow; >0.35 to 1 is red. a) F_ST_ calculated for each variant is indicated, with outlier variants coloured in red and non-outliers as light blue. b) F_ST_ calculated in windows inclusive of invariant sites. c) d_XY_ is calculated in windows inclusive of invariant sites. d) The values shown in subfigures b and c are subtracted to depict the difference between F_ST_ and d_XY_ values in each windowed region with the trendline in blue. e) Nucleotide diversity (π) is calculated in windows inclusive of invariant sites for *Bactrocera neohumeralis* with the trendline in blue. f) As above but showing *B. tryoni’s* nucleotide diversity. g) The recombination rate as predicted by ReLERNN for *B. neohumeralis* is indicated with grey shading showing the 95% confidence interval. h) As above but showing *B. tryoni’s* recombination rate.

**Figure 8.**
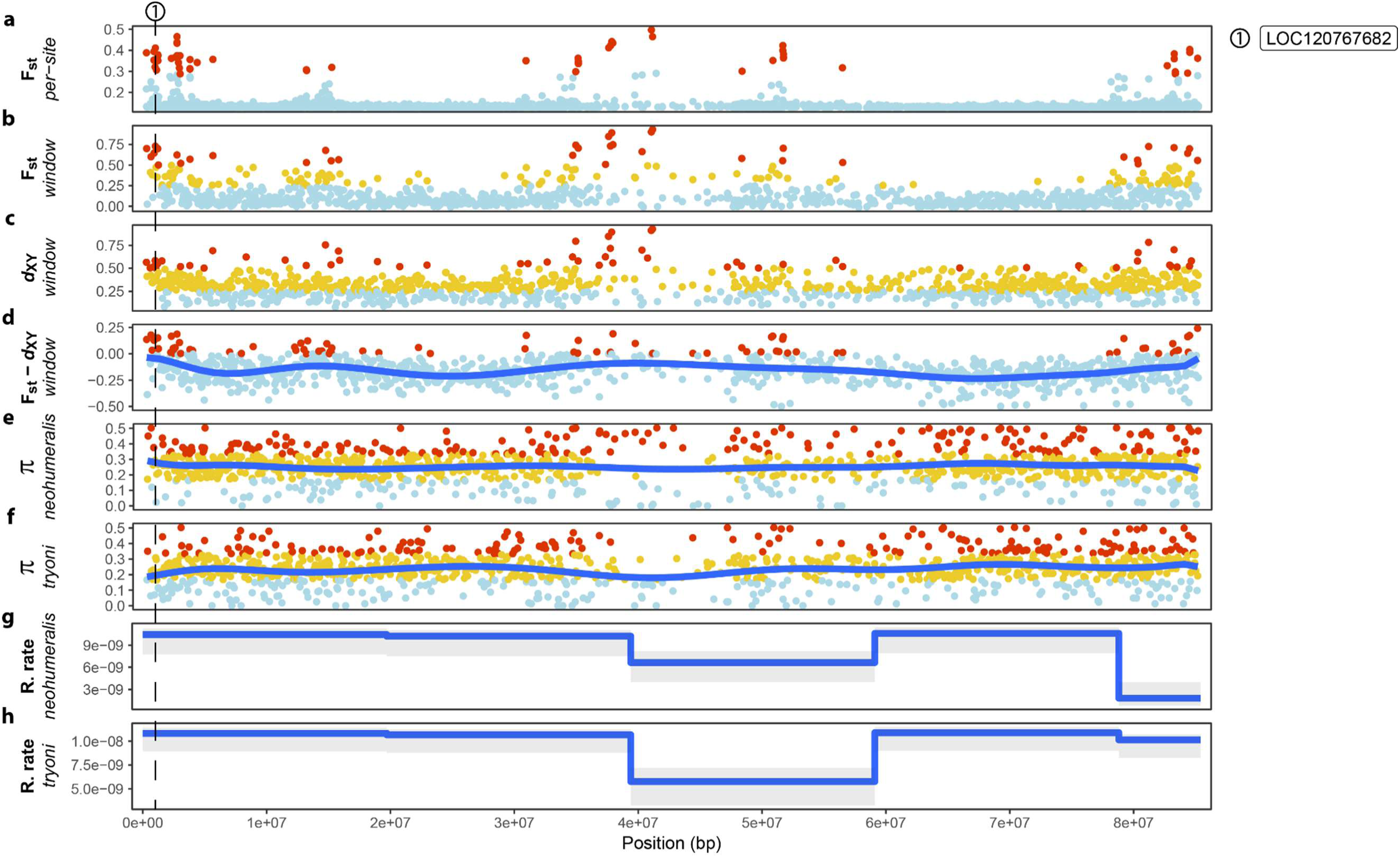
Population genetics statistics calculated for chromosome 2 labelled as NC_052500.1. Figure details are described in Fig. 7.

Mean nucleotide diversity estimates (π) produced similar results for each species (*B. tryoni* = 0.243962, *B. neohumeralis* = 0.242033). Nucleotide diversity estimates for each species were homogeneously distributed along the chromosomes with no significant clustering of elevated or lower rates observed (See Fig. 6 to Fig. 10). The LOSTRUCT results support the F_ST_ and d_XY_ values with significant separation between species along PC1.

**Figure 9.**
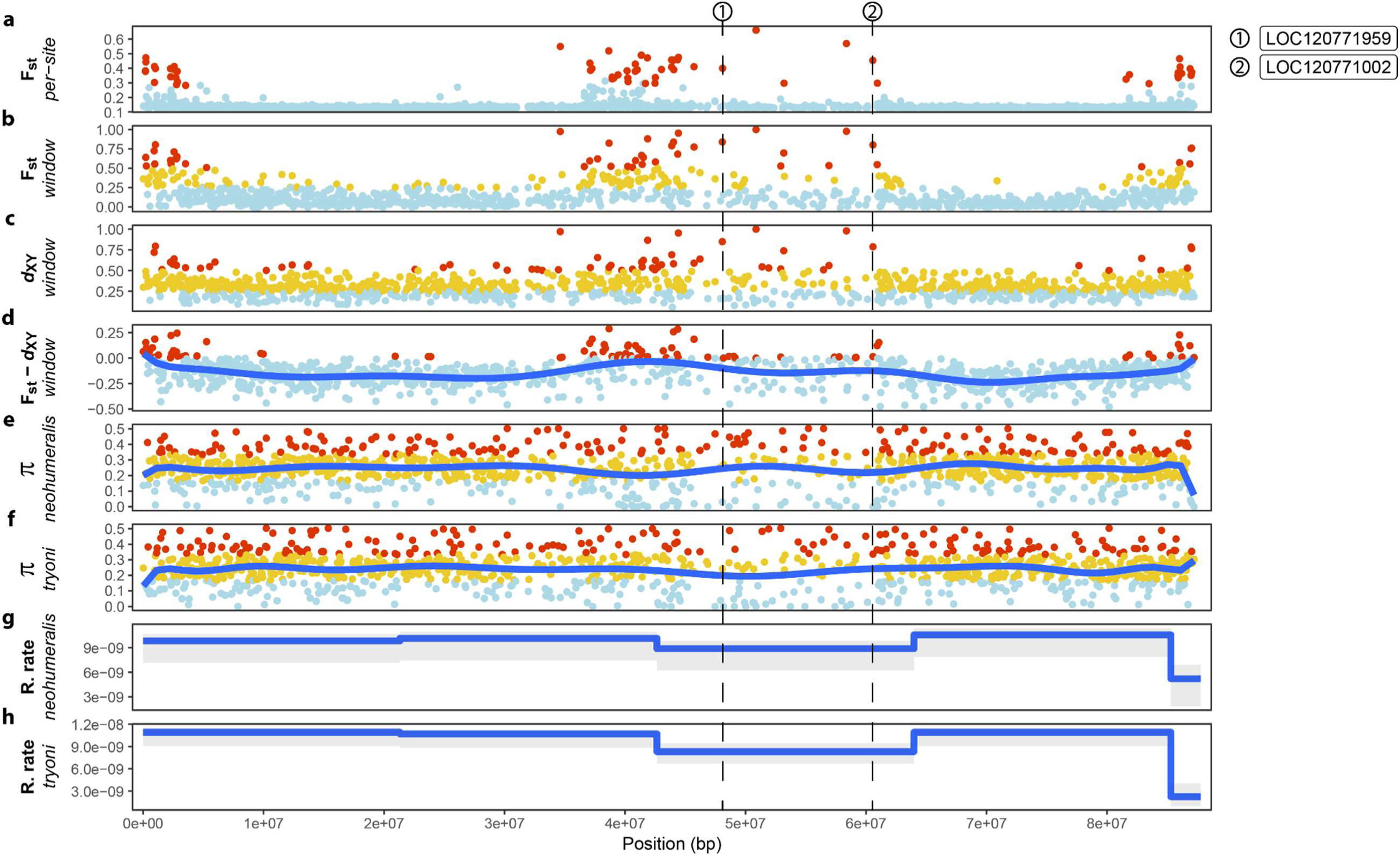
Population genetics statistics calculated for chromosome 3 labelled as NC_052501.1. Figure details are described in Fig. 7.

**Figure 10.**
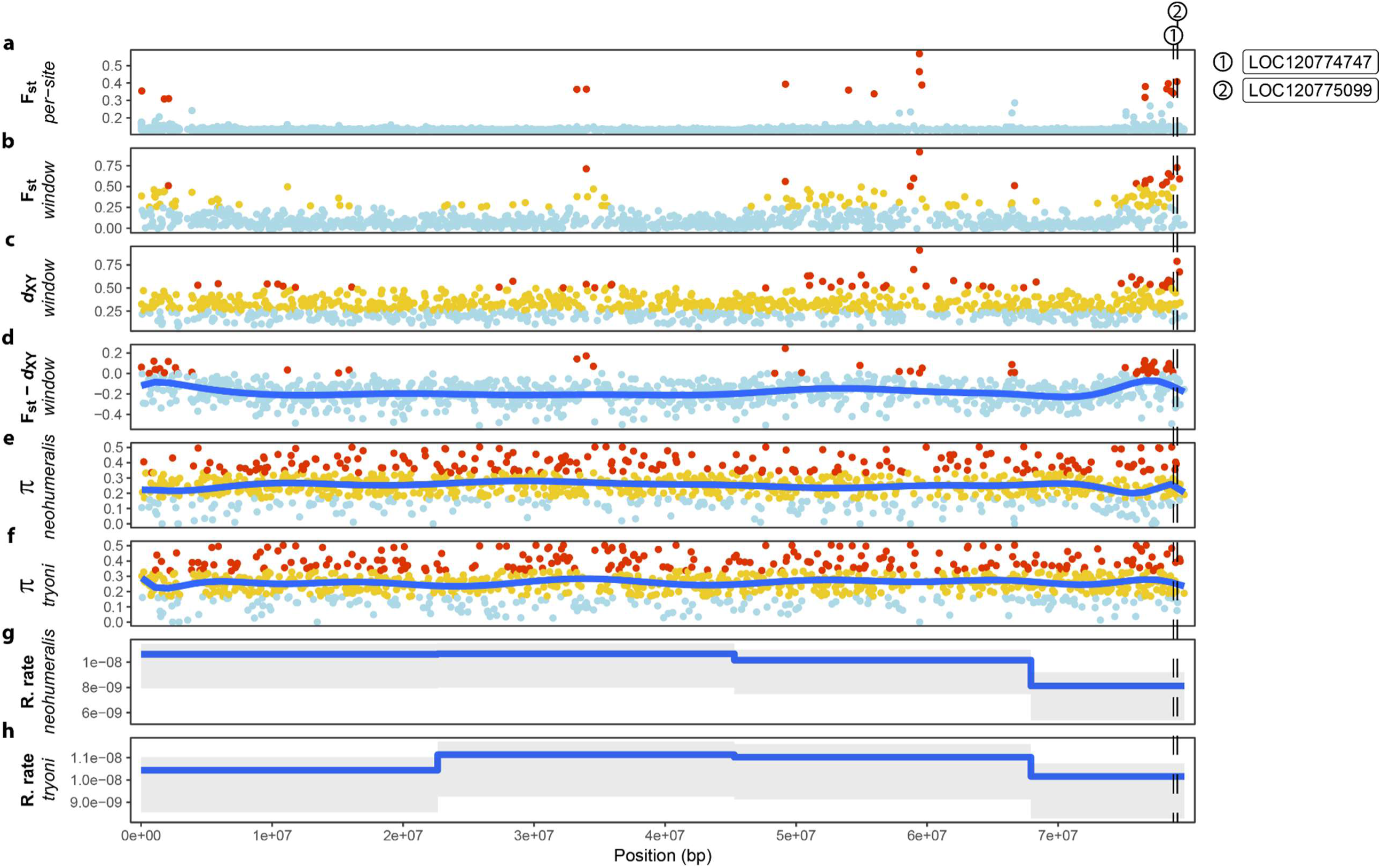
Population genetics statistics calculated for chromosome 3 labelled as NC_052502.1. Figure details are described in Fig. 7.

Overall, recombination rates within each species were low, with *B. tryoni* having a mean value of 9.02×10^-9^ per base and *B. neohumeralis* having a mean value of 9.78×10^-9^ per base. Recombination rates were overall consistently distributed along each chromosome with minor decreases observed towards the telomeres and the centromere with eight regions of decreased recombination rates occurring for each species (See Fig. 7 to Fig. 11). In each instance, these regions included clustered regions of elevated F_ST_ and d_XY_ values, however not all clustered regions of elevated F_ST_ and d_XY_ occurred in regions with lower recombination rates.

**Figure 11.**
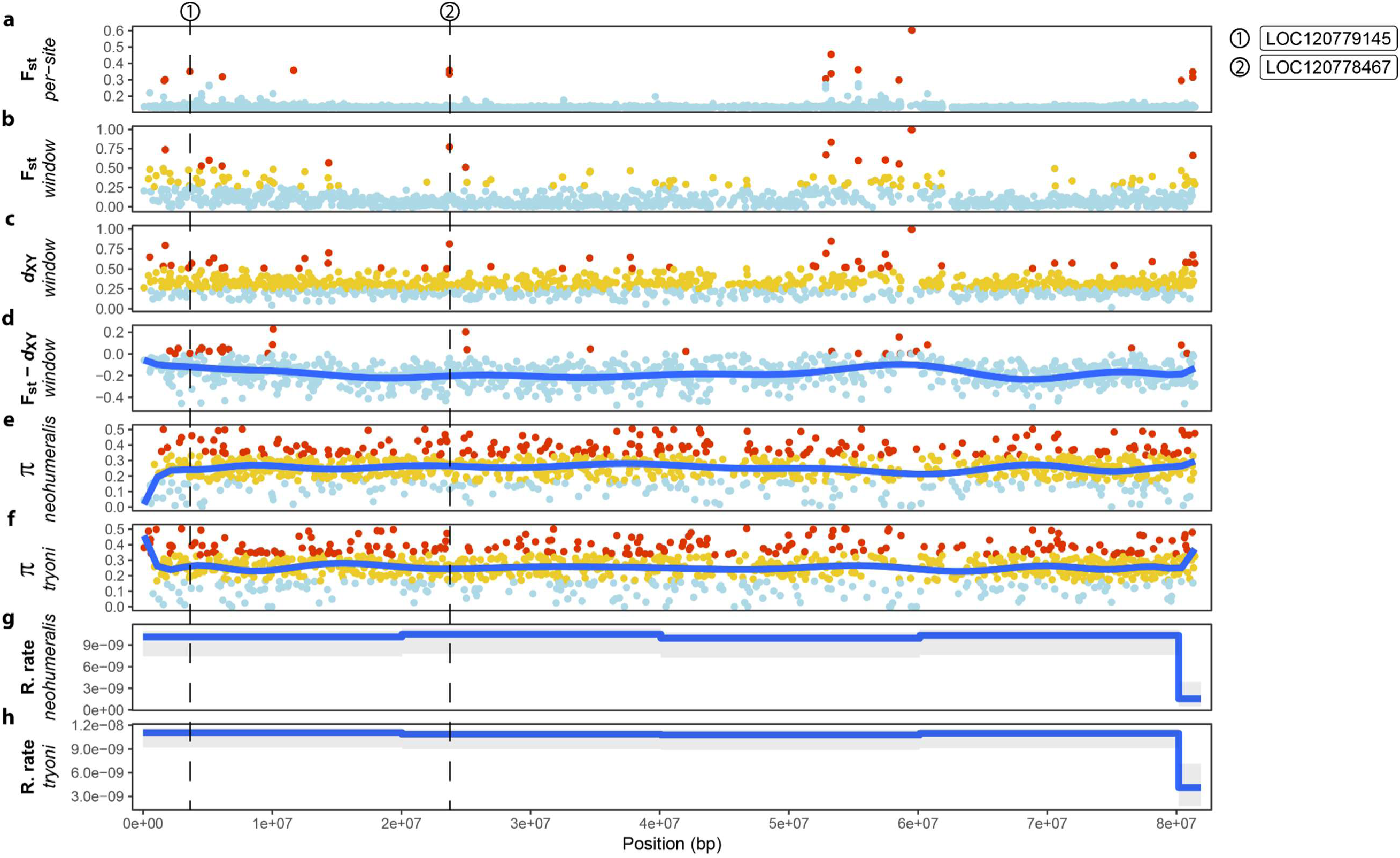
Population genetics statistics calculated for chromosome 3 labelled as NC_052503.1. Figure details are described in Fig. 7.

#### Outlier loci are clustered in specific chromosome segments between the species

Genes that contained outlier SNPs or were in close proximity to outlier SNPs were investigated, resulting in the identification of 266 genes. Enrichment of annotation terms indicates an overabundance of genes with the ‘abnormal neuroanatomy’ term (n=11 genes, p= 0.0164) within these candidate genes. Several neurology-related terms that were not enriched but still of interest included ‘nervous system development’, ‘central nervous system development’, ‘sensory system’, and ‘sensory organ development’. Additional non-enriched terms of interest included sleep and circadian rhythm-related terms, including: ‘abnormal sleep’, ‘circadian rhythm’, ‘abnormal circadian rhythm’, and ‘abnormal circadian behaviour’. Inositol-related terms were statistically enriched, that included ‘inositol biosynthetic process’ (n=1 gene, p= 0.0115) and ‘inositol-1,4,5-trisphosphate 3-kinase activity’ (n=1 gene, p= 0.0467). The 19 genes annotated with these terms are shown in Table S4.

### Ecological Data

#### No differences in seasonal activity between the species

Broad scale trap data collected over multiple years and sites throughout the sympatric range showed no difference in seasonal activity between *B. neohumeralis* and *B. tryoni* (Fig. 12 & Fig. S4). For any given sampling event there were often significantly more *B. tryoni* than *B. neohumeralis* trapped (See Table S5), but the abundance of both species over time were highly linearly correlated across nine sites with correlation values ranging from: r = 0.776 (Ayr, p < 0.001) to r = 0.993 (Maryborough, p < 0.001) (See Table S6). South Johnstone was the only site with a non-significant result (r = 0.926; p = 0.074) however, limited collections were recorded at this site (n = 4), providing insufficient statistical power to detect an association.

**Figure 12.**
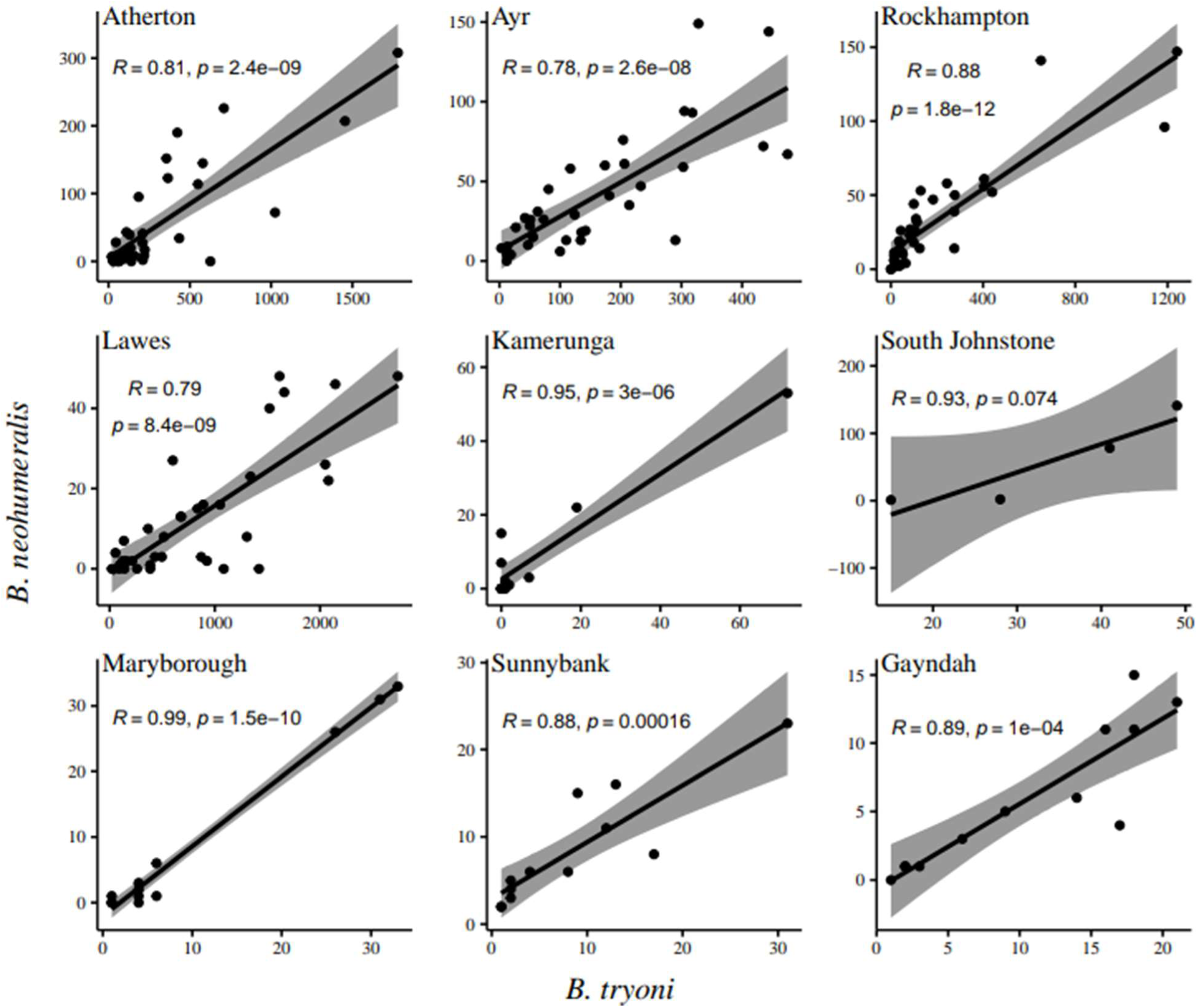
Correlation of seasonal abundance of *Bactrocera tryoni* and *B. neohumeralis* at multiple locations in Queensland, Australia. Scatterplots were developed using monthly mean of trap catches over 2 - 7 years. Shaded areas represent the 95% confidence intervals. Data from May (1961).

Trap data collected on a fine scale throughout the Wide Bay-Burnett region over a 12-month period shows that the abundance of both species across sites were significantly correlated over most collection periods with time, with the exception of the December period (r = 0.12, p = 0.57) where *B. neohumeralis* abundance decreased at a greater rate than *B. tryoni* after the October collection (Fig. 13 & Table S7). As observed in the broad scale analyses, there were often more *B. tryoni* than *B. neohumeralis* collected (See Table S8).

**Figure 13:**
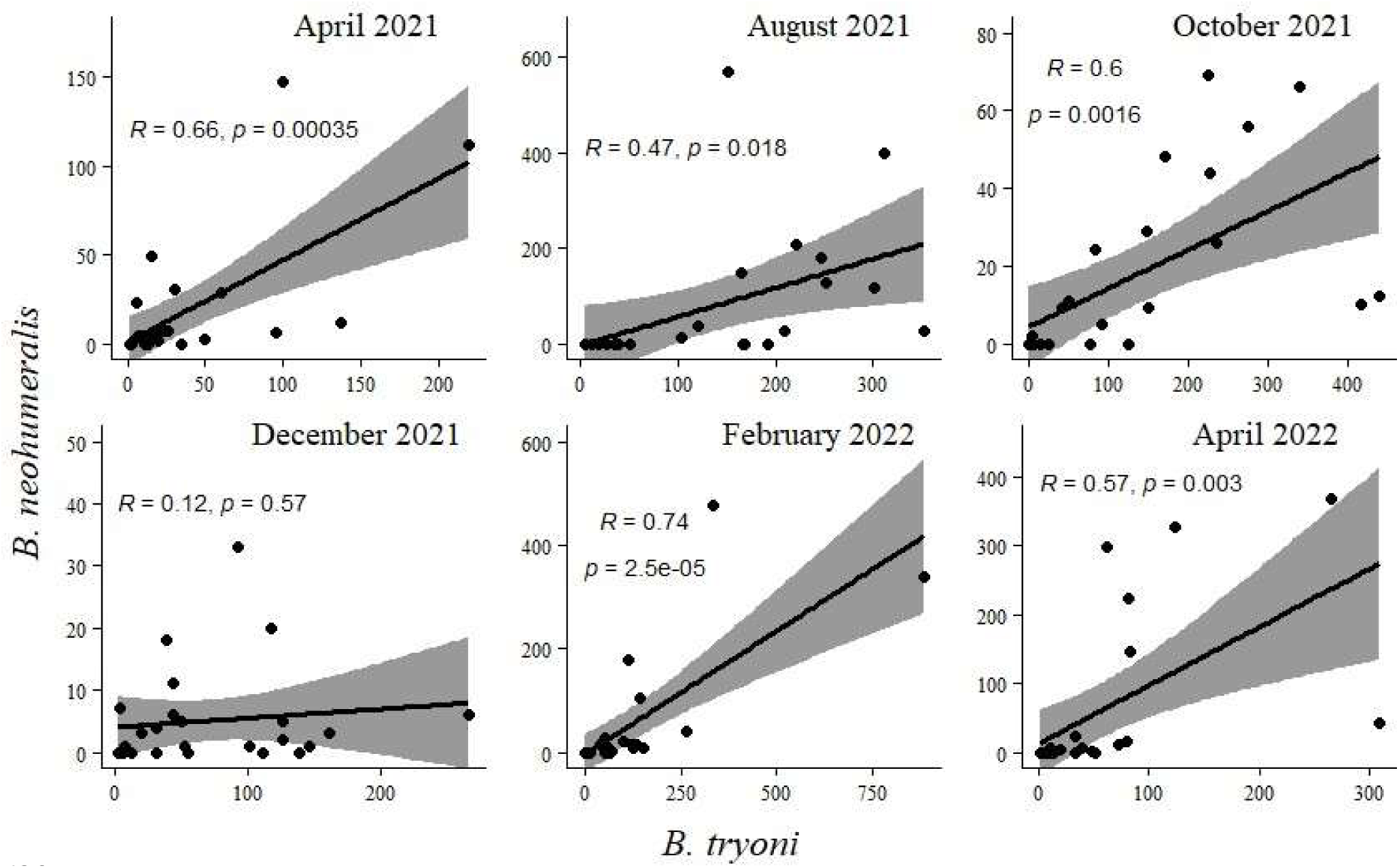
Correlation of fine scale seasonal abundance of *Bactrocera tryoni* and *B. neohumeralis* at multiple locations in the Wide Bay-Burnett region of Southeast Queensland. Shaded areas represent the 95% confidence intervals.

#### Correlated abundance across habitat types

There were significant positive correlations between *B. tryoni* and *B. neohumeralis* abundances in their habitat use (Fig. 14). Within a habitat type, there was a significant correlation in the two species monthly abundances for all habitat types, with the correlation ranging from r = 0.54 (dry forest, p <0.001) to r = 0.90 (mixed farming, p < 0.001) (See Table S10).

**Figure 14.**
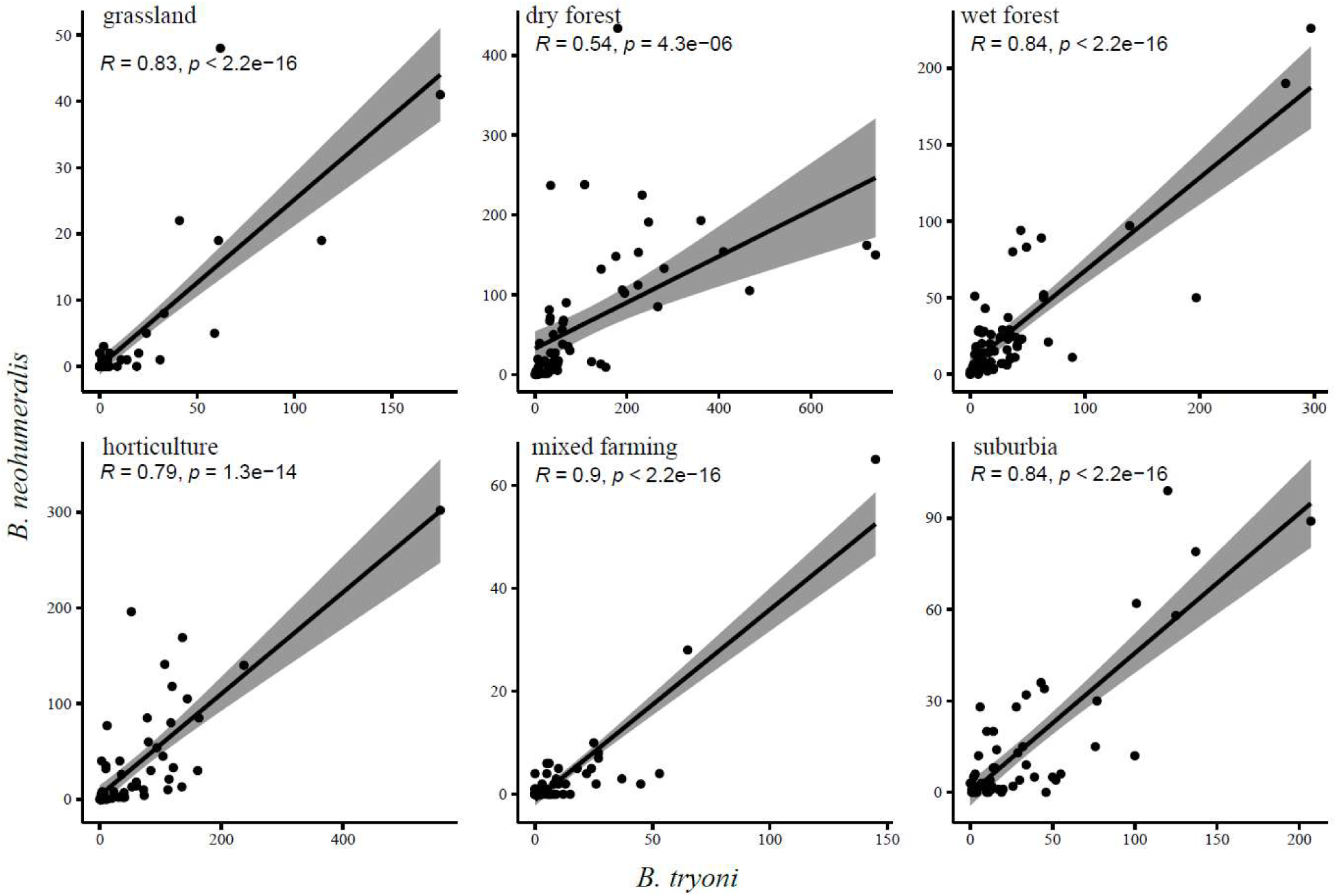
Correlation in habitat use of *Bactrocera tryoni* and *B. neohumeralis* in different human-defined habitat types within the Bundaberg district of Queensland. Each data point is the adult trap catch of both species for a trap run for three consecutive days. For each habitat type, nine traps were run per month for seven months. Shaded areas represent the 95% confidence intervals.

#### Landscape ecology

The LMM analysis included a total of 512 combinations of landscape predictors. Model averaging was performed using LMMs with ΔAICc < 2 from the top model (See Table S12). The top performing models retained eight predictors (See Table S13) from the nine included in the analysis, with only elevation not included in one of the top performing models. The top performing model was a combination of rainforest area, *Eucalyptus* woodland, latitude and longitude, with each predictor positively correlated with *B. neohumeralis* abundance. The 13 other top performing LMMs all included rainforest area as a predictor. No other predictor was included in all the top performing LMMs. Following model averaging only rainforest area was a statistically significant predictor of *B. neohumeralis* abundance (See Table S13) and was positively correlated. This pattern can be observed in Figure 15 (Ai, ii, v, vi, vii), where the high levels of abundance observed correlate with rainforests. In contrast, Ryan et al., (in review) found grazing area was the only significant predictor of *B. tryoni* abundance and was negatively correlated (See Fig. 15 B.viii, Table S14, and Table S15).

**Figure 15:**
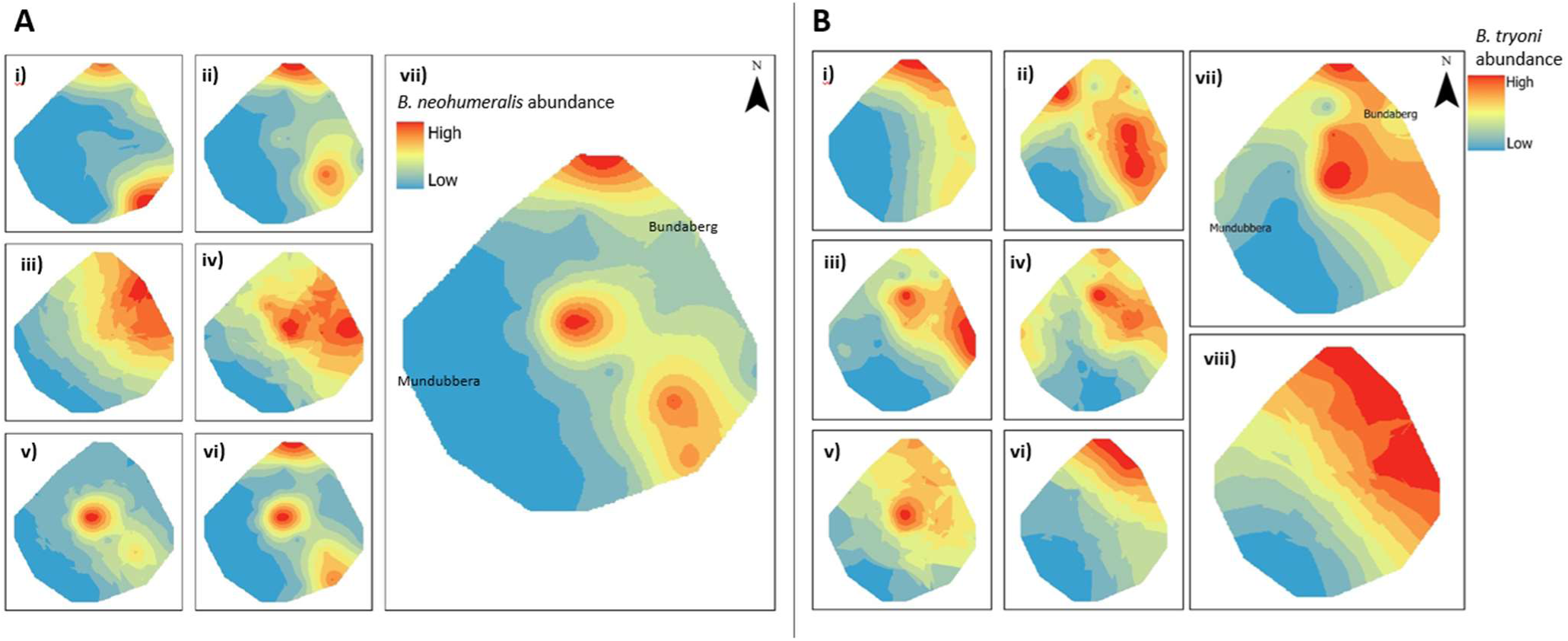
A) Empirical Bayesian kriging of *Bactrocera neohumeralis* abundance during each sampling period. i) April 2021, ii) August 2021, iii) October 2021, iv) December 2021, v) February 2022, vi) April 2022, vii) Average across all time periods. B) Empirical Bayesian kriging of *B. tryoni* abundance during each sampling period. i) April 2021, ii) August 2021, iii) October 2021, iv) December 2021, v) February 2022, vi) April 2022, vii) Average across all time periods, viii) Average abundance across all time periods excluding outlier from Good Night 2 in February 2022. Taken from Ryan et al. (In Review).

#### Limited differences in host use between species

A total of 1,607 fruits were sampled from the field and the emergence of *B. tryoni* and *B. neohumeralis* adults counted. For white sapote, mulberry, peach, plum, carambola and nectarine, sufficient samples were collected to carry out formal analyses. Highly significant correlations between the number of *B. tryoni* and *B. neohumeralis* adults reared from each of these fruit types were found (correlation ranging from r = 0.904, p <0.001 for white sapote to r = 0.952, p < 0.001 for peach) (Fig. 16 and Table S13). For guava, two collections occurred: a single *B. tryoni* and no *B. neohumeralis* emerged from the September 2013 collection while 1,609 *B. tryoni* and 1,353 *B. neohumeralis* emerged from the October 2013 collection; for feijoa a single collection occurred with equal numbers [71 flies each] of *B. tryoni* and *B. neohumeralis* emerging. For sapodilla, *Syzigium* spp., grumichama, hog plum, jabotica, white mulberry, avocado, black sapote, longan, mango and cashew, *B. tryoni* but no *B. neohumeralis* emerged.

**Figure 16.**
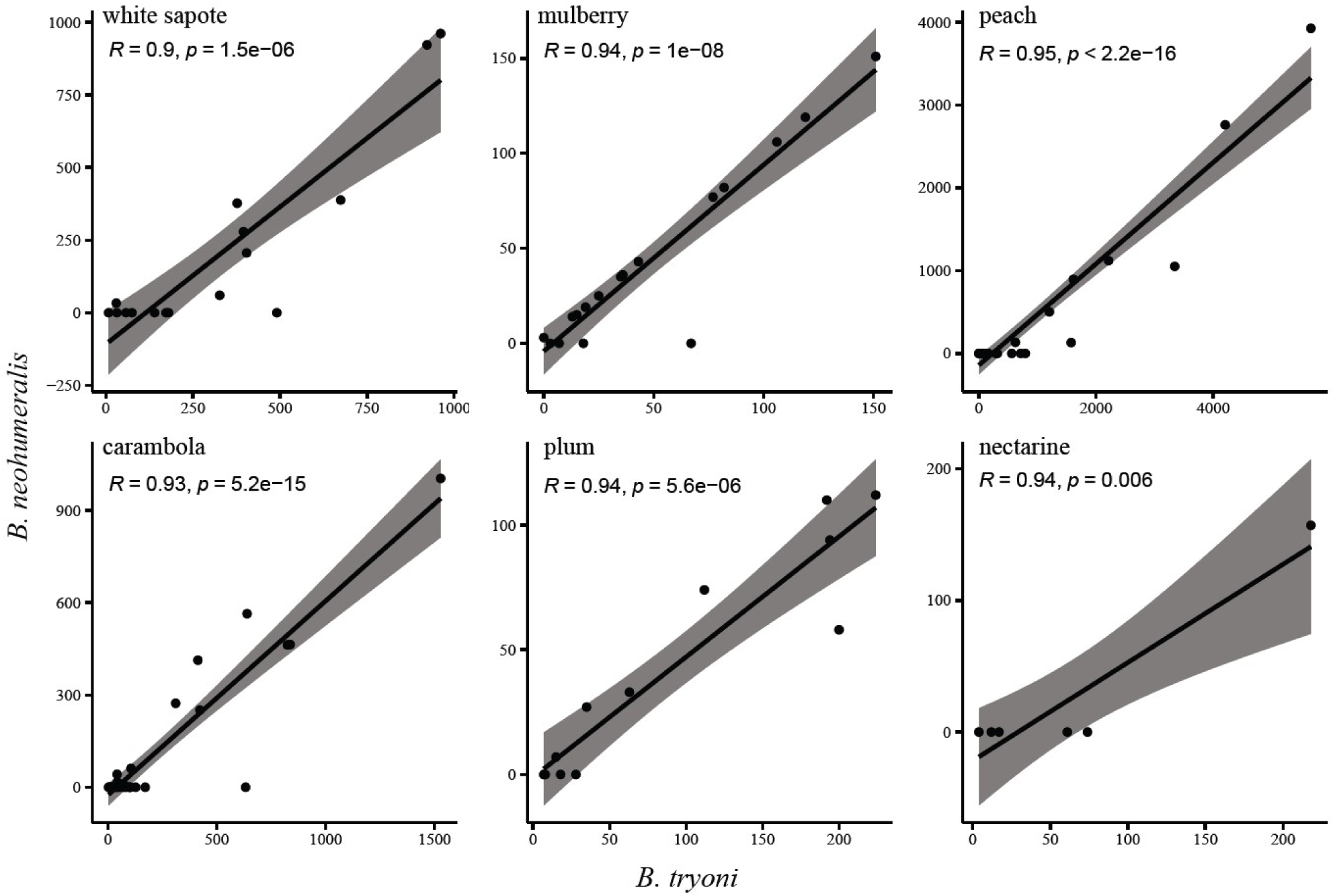
Correlation in host use of *Bactrocera tryoni* and *B. neohumeralis*. Each data point is the number of adult flies reared from an individual collection of multiple fruit pieces from a research orchard at Nambour, Queensland. Shaded areas represent the 95% confidence intervals.

## Discussion

### High nucleotide diversity suggests allopatric divergence through vicariance

Similar levels of nucleotide diversity (π) were estimated for *Bactrocera tryoni* (0.243962) and *B. neohumeralis* (0.242033). These values are consistent with prior studies comparing the nucleotide diversity of native (0.21552 – 0.22575) and invasive populations (0.19929 – 0.2199) of *B. tryoni* throughout Australia and the South Pacific (Parvizi et al., 2024) but substantially higher than those observed in other widespread *Bactrocera* species (*B. dorsalis* - 0.00148-0.0183) (Deschepper et al., 2023). High effective population sizes and high genetic diversity have previously been observed in population genetic studies on *B. tryoni* (Popa-Báez et al., 2020; Ryan et al., In Review) which may account for the elevated levels of nucleotide diversity observed in both species presented here. Low chromosome-wide recombination rates may also explain high nucleotide diversity as genetic mutations, typically purged through recombination (Kondrashov, 1994), are able to accumulate. Furthermore, low recombination rates may allow for the preservation of associations between linked alleles (Ortiz-Barrientos et al., 2016; Otto, 2009).

Yeap et al. (2020) observed a persistence in linkage between two *B. tryoni* - *B. neohumeralis* hybrid phenotypes involving humeral calli colour and time-of-day mating over 25 generations of hybrid backcrossing (without deliberate selection) despite a predicted linkage decay of 98.3% over 24 generations. The authors postulated that a reduction in recombination rates as a result of chromosomal rearrangements likely contributed to the persistence of linkage, they also suggested that an accumulation of nucleotide differences may also have been a factor. Low recombination rates have been observed in a number of Diptera (Cladera, 1981; Foster et al., 1980; Rössler, 1982) which can be attributed to an absence (or scarcity) of recombination in males (Zhao et al., 2003).

Consistently high levels of nucleotide diversity along each of the chromosomes coupled with predominantly low F_ST_ and d_XY_ values could be an indicator as to how initial divergence between these species occurred. Under a scenario of speciation with gene flow or allopatric speciation involving a small founder population, a genetic bottleneck is expected to reduce the nucleotide diversity of the population expanding into the novel environment (Coyne & Orr, 2004).

The low recombination rates observed in *B. tryoni* and *B. neohumeralis* may have allowed for the accumulation of high nucleotide diversity within each species post speciation, however, one would expect to see elevated F_ST_ and d_XY_ values across the chromosome as a result, which is not evident in our data. An alternative explanation is that a substantial vicariance event separated the ancestral species into two large populations, allowing speciation to occur in the absence of gene flow. During the period of geographic isolation, differential selection pressure would have led to the differences in the time-of-day mating between the species, limiting hybridization upon secondary contact (Coyne & Orr, 2004; Mayr, 1957).

### Genomic landscape indicates the timing of genetic divergence during speciation

A number of tightly clustered regions of elevated genetic differentiation appear to occur in regions with further reduced recombination along chromosomes 2-5 (See Fig. 8 to Fig. 11). These regions are consistent with models of hybridizing species where the linkage of genes involved in assortative mating with those associated with habitat preference and performance is considered necessary for speciation with gene flow (Felsenstein, 1981; Ortíz-Barrientos et al., 2002; Yeaman, 2013). The formation of the tightly clustered regions of genetic differentiation observed in data presented here may have been associated with the initial divergence of the two species if speciation occurred in sympatry. However, based on the high levels of nucleotide diversity observed it is perhaps more likely that these tightly clustered regions were important in maintaining species boundaries during the early periods of secondary contact where the species diverged in allopatry. If reproductive barriers were porous during secondary contact, the strong linkage of locally adapted alleles and genes associated with assortative mating would have likely strengthened the reproductive barriers between the two species, limiting the homogenizing effect of hybridization (Felsenstein, 1981; Ortíz-Barrientos et al., 2002).

In addition to those clustered regions with reduced rates of recombination, a number of regions of elevated genetic diversity without a reduction in recombination rates are present. Elevated genetic diversity without reduced rates of recombination are consistent with the patterns expected under the model of divergence after speciation where populations experience a period of geographic isolation allowing divergence to occur in the absence of gene flow (Cruickshank & Hahn, 2014). Under this model, differential selective pressures drive divergence between the species through selective sweeps and the absence of gene flow allows them to persist without a reduction in recombination. The level of absolute genetic divergence (d_XY_) in relation to relative genetic differentiation (F_ST_) could provide insight into the timing of these selective sweeps relative to the completion of speciation between taxa. Where both statistics are elevated, the selective sweep likely occurred soon after speciation. In contrast, regions with elevated F_ST_ and non-elevated d_XY_ could suggest a more recent selective sweep (Cruickshank & Hahn, 2014; Ravinet et al., 2017).

### Assortative mating maintains species boundaries in sympatry

Genetic comparisons of *B. tryoni* and *B. neohumeralis* individuals collected throughout the overlapping range showed no evidence of hybridization between the species. This was despite samples of each species regularly being collected from the same traps, i.e., in immediate sympatry. Several intermediate phenotypes were collected, however, these samples presented strongly as *B. neohumeralis* genotypes with no indication of hybridization. This is consistent with the findings of prior studies of the species pair and supports the theory that strong reproductive barriers exist in natural conditions (Gilchrist & Ling, 2006; Pike, 2004; Yeap et al., 2020). The lack of evidence of hybridization identified from natural populations in this study is consistent with the results of a recent diagnostic tool development study where no hybrids were identified from *B. tryoni* (n = 430) and *B. neohumeralis* (n = 99) samples collected across > 40 collection sites throughout their overlapping range (Starkie et al., 2023). These results along with asymmetric hybridization observed in a prior laboratory-based study (Yeap et al., 2020), suggest that assortative mating is the most prevalent form of reproductive isolation responsible for maintaining species boundaries between the pair. However, the existence of hybrid selection in natural environments cannot be discounted, definitively.

While genes related to mating behaviour and circadian rhythm were not enriched in the gene enrichment analysis, 19 candidate genes were located in the tightly clustered regions of high genetic differentiation. These genes are postulated as being responsible for maintaining separation between the two species and forming the time-of-day premating reproductive barrier through assortative mating. Further analysis is required to determine whether these putative genes associated with assortative mating are linked to locally adapted genes, given the apparent lack of ecological niche differentiation between the species as discussed below.

### Lack of competitive displacement allows co-existence

Competitive displacement has long been suggested as an important mechanism facilitating divergence and speciation (Rice & Salt, 1988; Thoday & Gibson, 1962; Via, 2001). The ability of two nascent species to co-exist by minimizing competition, or by utilizing non-competitive ecological space, should be considered central to early species divergence and persistence (Germain et al., 2021). Where two herbivore populations share resources, then displacement can occur in time (i.e. seasonal activity), location, or host usage (Clarke & Measham, 2022; Facon et al., 2021). However, the data presented here show no evidence for this in the *B. tryoni*/*B. neohumeralis* sibling pair. Both species were active at the same time of the year and in the same habitats within a landscape. *Bactrocera tryoni* was recovered from a greater range of host fruits than *B. neohumeralis* and that may be seen as evidence for competitive displacement (i.e. competitive advantage by *B. neohumeralis* resulting in *B. tryoni* adapting to novel hosts), but it is more likely that this pattern simply reflects *B. tryoni*’s greater abundance in the environment and so greater likelihood of being sampled, rather than a competition avoidance mechanism. This is because when both species were reared from the same fruit, their numbers were closely correlated. In cases where competition between fruit flies has been known to cause changes in field host use patterns, one species will dominate in a given host, pushing a second species to host fruits that the first only irregularly uses (Charlery de la Masselière et al., 2017; Hassani et al., 2022); this pattern was not observed in these data. The positive correlation in the number of adults of both species from fruit also offers field data supporting the work of Kay et al. (2024), who in laboratory studies found no interspecific larval competition between the species.

Ecological traits typically associated with competition driven divergence between fruit fly species (i.e. differential host use, habitat usage, and seasonal abundance (Clarke & Measham, 2022; Duyck et al., 2004; Inskeep et al., 2021)) do not appear to differ between taxa in this study, however, the results of our LMM and those of Ryan et al. (In Review) indicate there may be some minor difference in habitat preference. We show that *B. neohumeralis* appears to prefer rainforest systems which is a subset of the habitat preference of *B. tyroni* which is only affected by the presence of cleared grazing area. Therefore, the difference in mating time remains the only known distinction between the species pair.

## Author contributions

A.R.C. and P.P. conceived the study. M.I., N.K., C.G.M., J.R., S.B. and, B.M. conducted field work and collected samples for molecular and ecological analysis. M.I., N.K., J.R., A.R.C. and, P.P. designed, collated, and analyzed ecological data. Z.S. and P.P. designed and performed bioinformatics and SNP analysis. M.I., Z.S., P.P. collated analyzed molecular data. M.I., Z.S., N.K., J.R., C.G.M., M.S., A.R.C., D.H. and P.P. contributed to the writing of the manuscript. All authors have read and approved the final version.

## Supporting information

Supplementary Section

Supplementary Section 2

## Acknowledgements

We thank E. Vueti who carried out much of the Bundaberg sampling as part of an uncompleted postgraduate study, and L. Senior (QDAF) for her assistance with rearing flies from fruit that provided data for the host utilization study. We also thank The Southern Downs Regional Council, The Granite Belt Growers Association, B. Doohan, The Greenhoose, Lockhart River, D. Masters, J. Kerr and J. Whistler for their assistance in collecting samples for the molecular analysis.

This research was partially funded by the ARC ITTC project IC150100026, Centre for Fruit Fly Biosecurity Innovation awarded to A.R.C. and P.P. During the final stages of writing P.J.P., D.H., M.S., B.M., and S.B. received support through Fresh and Secure Trade Alliance. The Fresh and Secure Trade Alliance is funded through the Hort Frontiers International Markets Fund, part of the Hort Frontiers strategic partnership initiative developed by Hort Innovation, with co-investment from the Department of Agriculture and Fisheries, Department of Primary Industries and Regional Development, Department of Energy,

Environment and Climate Action, Department of Tourism, Industry and Trade, Department of Primary Industries and Regions, Department of Natural Resources and Environment, Queensland University of Technology, James Cook University, Western Sydney University, Australian Blueberry Growers’ Association, GreenSkinAvocados, and contributions from the Australian Government and the strawberry and avocado R&D levy.

The authors declare no conflicts of interest.

## Information on Data Deposition

BioProject PRJNA1008389, Zachary Stewart, 2023, DArTseq of *Bactrocera neohumeralis* and *tryoni* (Queensland fruit fly / Qfly), NCBI SRA, BioProject PRJNA1008389.

