## Supplementary Section for "Strong genetic differentiation but limited niche partitioning in a sympatric species pair separated by an allochronic reproductive barrier"

**Supplementary Data**

**Table S1.** Trapping records from 19 collection sites for molecular data analysis. The individuals with an intermediate phenotype were allocated to a separate “site” labelled intermediate (acronym – I) for the fastSTRUCTURE analysis.

| **Collection site** | **Site acronym** | **Latitude** | **Longitude** | ***B. neohumeralis*** | ***B. tryoni*** | **Intermediate phenotype** |
| --- | --- | --- | --- | --- | --- | --- |
| Bowen | BO | -20.01210 | 148.24627 | 5 | 6 |  |
| Brisbane East | BRI_E | -27.49136 | 153.04577 | 4 | 6 |  |
| Bundaberg | BU | -24.85048 | 152.35064 | 6 | 6 |  |
| Cleveland | CLV | -27.52700 | 153.27325 | 6 | 6 |  |
| Cooktown | CK | -15.47581 | 145.24709 | 6 | 0 |  |
| Coomera | COO | -27.86239 | 153.32151 | 6 | 6 |  |
| Fordsdale | FD | -27.71585 | 152.09481 | 3 | 6 |  |
| Gold Coast | GC | -28.01666 | 153.43600 | 6 | 6 |  |
| Hervey Bay | HB | -25.30250 | 152.88861 | 6 | 6 |  |
| Iron Range | IR | -12.76415 | 143.28570 | 10 | 0 |  |
| Mackay | MA | -21.15106 | 149.19892 | 6 | 0 |  |
| Moreton | MT | -12.45357 | 142.63792 | 5 | 6 |  |
| Mt Nathan | MtN | -27.99456 | 153.27574 | 6 | 1 | 3 |
| Noosaville | NS | -26.40220 | 153.05400 | 6 | 6 |  |
| North Ipswich | NI | -27.60649 | 152.76286 | 4 | 6 |  |
| Rockhampton | RH | -23.31264 | 150.51791 | 9 | 5 |  |
| Scarborough | Sca | -27.21079 | 153.11375 | 6 | 6 |  |
| Stanthorpe | STN | -28.73358 | 151.98922 | 0 | 3 |  |
| Townsville | TV | -19.22269 | 146.67944 | 2 | 0 |  |

**Table S2.** See Supplementary Section 2.

**Table S3.** See Supplementary Section 2.

**Table S4.** See Supplementary Section 2.

**Table S5**. Trap catches of *B. tryoni*: *B. neohumeralis* for seasonal activity in various locations in Queensland, Australia. For Atherton, Ayr, Rockhampton and Lawes, weekly data were summed to a calendar month. For Maryborough, Sunnybank, Kamerunga and Gayndah, percent mean of total trap catches was calculated. For South Johnstone, mean seasonal data (for winter, summer, autumn, and spring seasons) only, were available. For Rita Island, Nambour, Withcott, Toowoomba, Offham and Stanthorpe, only total trap catch data were available.

| **Stations** | **Years trap maintained** | ***B. tryoni* per trap per year** | ***B. neohumeralis* per trap per year** | ***B. tryoni: B. neohumeralis*** |
| --- | --- | --- | --- | --- |
| Kamerunga | 4 | 48.6 | 14 | 3.47 : 1 |
| Atherton | 4 | 301.9 | 44.6 | 6.77 : 1 |
| South Johnston | 3 | 4.4 | 7.4 | 0.59 : 1 |
| Rita island  Ayr | 3  4 | 37.6  160.5 | 25.1  38.7 | 1.5 : 1  4.13 : 1 |
| Rockhampton | 3 | 222.1 | 36.8 | 6.04 : 1 |
| Maryborough  Gayndah | 3  2 | 34.97  550.1 | 7.9  6.6 | 4.43 : 1  83.35 : 1 |
| Nambour | 1 | 148.1 | 15.1 | 9.81 : 1 |
| Sunnybank | 3 | 734.1 | 59 | 25.31 : 1 |
| Lawes  Withcott  Toowoomba  Offham | 6  2  6  1 | 1065.3  231.2  313.98  39.2 | 14.2  2.6  3.34  0 | 75.02 : 1  88.92 : 1  94.01 : 1  - |
| Stanthorpe | 4 | 191.7 | 0.125 | 1533.6 : 1 |

**Table S6.** Results for the broad scale seasonal abundance data correlation analysis between *B. tryoni* and *B. neohumeralis*.

| **Location** | **t-statistic** | **df** | **p-value** | **Confidence intervals** | **Pearson’s correlation coefficient** |
| --- | --- | --- | --- | --- | --- |
| Atherton | 8.034 | 34 | <0.001 | 0.654 - 0.900 | 0.810 |
| Ayr | 7.185 | 34 | <0.001 | 0.601 - 0.880 | 0.776 |
| Gayndah | 6.198 | 10 | <0.001 | 0.648 - 0.969 | 0.891 |
| Kamerunga | 9.325 | 10 | <0.001 | 0.817 - 0.985 | 0.947 |
| Lawes | 7.578 | 34 | <0.001 | 0.627 - 0.889 | 0.793 |
| Maryborough | 26.219 | 10 | <0.001 | 0.974 - 0.998 | 0.993 |
| Rockhampton | 10.747 | 34 | <0.001 | 0.774 - 0.937 | 0.879 |
| South Johnstone | 3.459 | 2 | 0.074 | -0.321 - 0.998 | 0.926 |
| Sunnybank | 5.837 | 10 | <0.001 | 0.616 - 0.966 | 0.879 |

**Table S7:** Results for the fine scale seasonal abundance data correlation analysis between *B. tryoni* and *B. neohumeralis.*

| **Location** | **t-statistic** | **df** | **p-value** | **Confidence intervals** | **Pearson’s correlation coefficient** |
| --- | --- | --- | --- | --- | --- |
| April 2021 | 4.186 | 23 | <0.001 | 0.355 - 0.836 | 0.657 |
| August 2021 | 2.539 | 23 | 0.018 | 0.089 - 0.728 | 0.468 |
| October 2021 | 3.567 | 23 | 0.001 | 0.264 - 0.803 | 0.597 |
| December 2021 | 0.582 | 23 | 0.566 | -0.288 - 0.492 | 0.120 |
| February 2022 | 5.254 | 23 | <0.001 | 0.485 - 0.878 | 0.739 |
| April 2022 | 3.318 | 23 | 0.003 | 0.224 - 0.787 | 0.569 |

**Table S8.** See Supplementary Section 2.

**Table S9.** See Supplementary Section 2.

**Table S10.** Results for the habitat use data correlation analysis between *B. tryoni* and *B. neohumeralis*.

| **Location** | **t-statistic** | **df** | **p-value** | **Confidence intervals** | **Pearson’s correlation coefficient** |
| --- | --- | --- | --- | --- | --- |
| grassland | 11.918 | 66 | <0.001 | 0.732 - 0.889 | 0.826 |
| dry forest | 5.047 | 61 | <0.001 | 0.341 - 0.697 | 0.543 |
| wet forest | 12.062 | 60 | <0.001 | 0.749 - 0.902 | 0.841 |
| horticulture | 10.085 | 61 | <0.001 | 0.675 - 0.868 | 0.791 |
| mixed farming | 16.364 | 61 | <0.001 | 0.843 - 0.940 | 0.902 |
| suburbia | 11.555 | 56 | <0.001 | 0.742 - 0.902 | 0.836 |

**Table S11.** Model support for top performing linear mixed models (LMMs) correlating *Bactrocera neohumeralis* abundance with landscape predictors after model averaging (top performing models all have ∆AICc < 2 prior to model averaging). Models are ranked according to AIC corrected for finite sample size (∆AICc) and Akaike weight (ω). Degrees of freedom (df) for each model are also given.

| **Model** | **∆AICc** | **ω** | **df** |
| --- | --- | --- | --- |
| Rainforest + *Eucalyptus* + Latitude + Longitude | 0 | 0.12 | 7 |
| Rainforest + *Eucalyptus* + Watersource | 0.24 | 0.11 | 6 |
| Rainforest + *Eucalyptus* + Watersource + Produce | 0.27 | 0.1 | 7 |
| Rainforest + Watersource + Grazing + Residential | 0.6 | 0.09 | 7 |
| Rainforest + *Eucalyptus* + Latitude + Longitude + Produce | 0.74 | 0.08 | 8 |
| Rainforest + Latitude + Longitude + Grazing + Produce | 0.96 | 0.07 | 7 |
| Rainforest + *Eucalyptus* + Watersource + Latitude | 0.97 | 0.07 | 6 |
| Rainforest + *Eucalyptus* + Latitude | 1.13 | 0.07 | 8 |
| Rainforest + *Eucalyptus* + Latitude + Longitude + Residential | 1.15 | 0.07 | 6 |
| Rainforest + *Eucalyptus* + Longitude | 1.24 | 0.06 | 7 |
| Rainforest + *Eucalyptus* + Longitude + Produce | 1.54 | 0.06 | 8 |
| Rainforest + *Eucalyptus* + Watersource + Latitude + Produce | 1.73 | 0.05 | 7 |
| Rainforest + Latitude + Longitude + Residential | 1.83 | 0.05 | 7 |
| Rainforest + Longitude + Grazing + Residential | 7.48 | 0 | 8 |

**Table 12.** Statistical support for each landscape predictor following model averaging. Reported are the linear coefficient, standard error (Std. error), Z-score, p value, 2.5% and 97.5% confidence intervals (CI). Bold text denotes statistical significance.

| **Variable** | **Coefficient** | **Std. error** | **Z-value** | **p-value** | **2.5% CI** | **97.5% CI** |
| --- | --- | --- | --- | --- | --- | --- |
| (Intercept) | -2017 | 2928 | 0.69 | 0.49 | -7772.46 | 3739.06 |
| *Eucalyptus* | 0.63 | 0.42 | 1.48 | 0.14 | 0.13 | 1.42 |
| Rainforest | 2.05 | 0.45 | 4.51 | **6.4x10^-6^** | 1.16 | 2.94 |
| Latitude | 14.86 | 18.92 | 0.78 | 0.43 | -5.83 | 62.57 |
| Longitude | 15.91 | 19.58 | 0.81 | 0.42 | -0.16 | 63.74 |
| Watersource | -0.05 | 0.08 | 0.72 | 0.47 | -0.25 | 0.00 |
| Produce | 0.17 | 0.34 | 0.50 | 0.62 | -0.25 | 1.37 |
| Grazing | -0.10 | 0.26 | 0.36 | 0.72 | -1.30 | -0.09 |
| Residential | -0.16 | 0.34 | 0.48 | 0.63 | -1.39 | 0.11 |

**Table S13.** Results for the host use data correlation analysis between *B. tryoni* and *B. neohumeralis*.

| **Location** | **t-statistic** | **df** | **p-value** | **Confidence intervals** | **Pearson’s correlation coefficient** |
| --- | --- | --- | --- | --- | --- |
| white sapote | 7.9493 | 14 | <0.001 | 0.742 – 0.967 | 0.905 |
| mulberry | 10.749 | 16 | <0.001 | 0.836 – 0.977 | 0.937 |
| peach | 17.047 | 30 | <0.001 | 0.903 – 0.977 | 0.952 |
| carambola | 13.792 | 32 | <0.001 | 0.854 – 0.962 | 0.983 |
| plum | 8.705 | 10 | <0.001 | 0.795 – 0.983 | 0.940 |
| nectarine | 5.322 | 4 | 0.006 | 0.518 – 0.993 | 0.936 |

**Table S14.** Top five performing linear mixed models (LMMs) correlating *B. tryoni* abundance with landscape predictors. Models are ranked according to AIC corrected for finite sample size (∆AICc) and Akaike weight (ω). Degrees of freedom (df) for each model are also given. Taken from Ryan et al., (in review).

| Model | ∆AICc | ω | df |
| --- | --- | --- | --- |
| Grazing + Watersource | 0.00 | 0.22 | 5 |
| Grazing | 0.01 | 0.22 | 4 |
| Grazing + Latitude | 0.45 | 0.18 | 5 |
| Grazing + Longitude | 1.40 | 0.11 | 5 |
| Grazing + Watersource + Latitude | 1.56 | 0.10 | 6 |
| Grazing + Latitude + Longitude | 1.82 | 0.09 | 6 |
| Grazing + Watersource + Elevation | 1.91 | 0.08 | 6 |

**Table S15.** Results of model averaging for the top five performing linear mixed models (LMMs). Reported are the linear coefficient, standard error (Std. error), Z-score, p value, 2.5% and 97.5% confidence intervals (CI). Bold text denotes statistical significance in a model and CIs which do not overlap 0. Taken from Ryan et al., (in review).

| Variable | Coefficient | Std. error | Z-value | p value | 2.5% CI | 97.5% CI |
| --- | --- | --- | --- | --- | --- | --- |
| (Intercept) | -31.32 | 247.3 | 0.125 | 0.900 | -520.9 | 458.3 |
| Grazing | -0.132 | 0.388 | 3.352 | **8x10^-4^** | **-0.209** | **-0.055** |
| Watersource | -0.006 | 0.010 | 0.608 | 0.543 | -0.035 | 0.006 |
| Latitude | 0.126 | 2.384 | 0.525 | 0.600 | -2.181 | 9.082 |
| Longitude | 0.523 | 1.604 | 0.323 | 0.747 | -2.775 | 8.063 |
| Elevation | 8x10^-4^ | 0.005 | 0.163 | 0.870 | -0.019 | 0.038 |

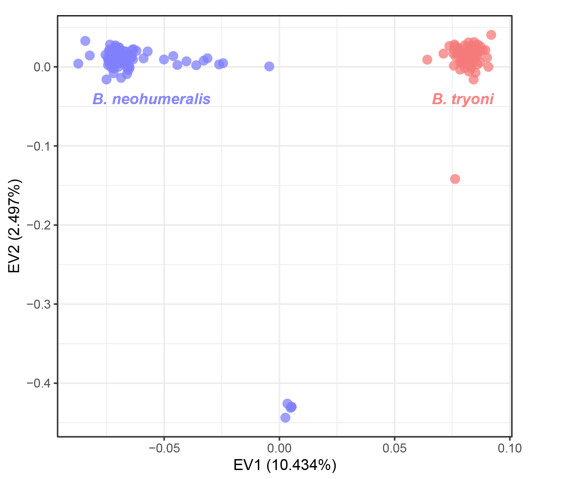

**Figure S1.** Principal component analysis (PCA) of single nucleotide polymorphisms (SNPs) *Bactrocera neohumeralis* and *B.* *tryoni*, including outlier samples, with the values from the first two eigenvectors plotted for each sample. Eigenvectors one (horizontal) and two (vertical) explain 10.434% and 2.497% of the variance in the data, respectively.

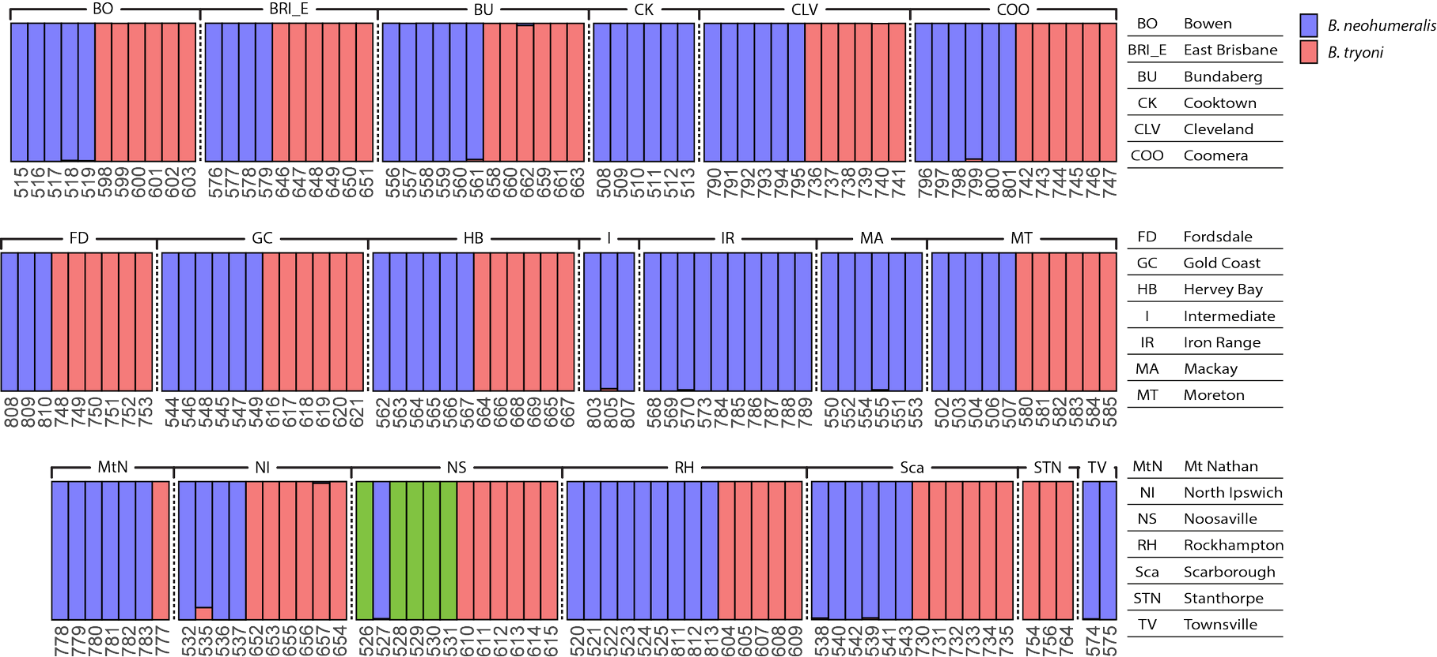

**Figure S2.** fastSTRUCTURE results at cluster size (K) equals three for single nucleotide polymorphisms (SNPs) for *Bactrocera neohumeralis* and *B. tryoni*. Red, blue, and green colouration of the bars indicates the inferred population admixture proportions as a percentage (numbers not shown). Three-digit numbers along the bottoms of each row of bars are the genotype ID codes.

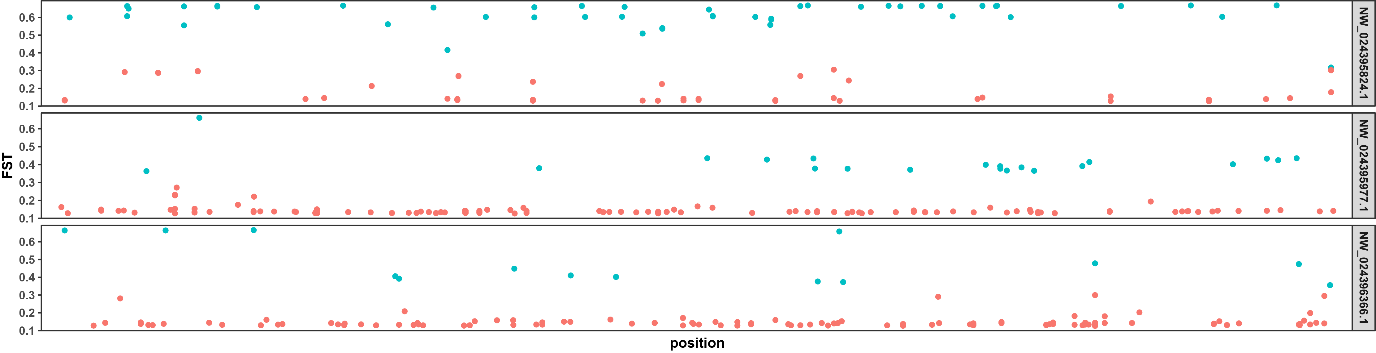

**Figure S3**. Unplaced contigs positioned and scaled along the X-axis in proportion to the five autosomal chromosomes displayed in Fig. 4. Fst values for each contig were computed and are plotted as the Y-axis value (left), with their NCBI sequence identifier shown to the right. Outlier SNPs are highlighted in blue.

Mean number of flies

|  | B. neohumeralis | B. tryoni |
| --- | --- | --- |
| Jan | 61 | 223.2 |
| Feb | 97.6667 | 129.4 |
| Mar | 65.3333 | 181.8 |
| Apr | 58.3333 | 122.4 |
| May | 23 | 72.25 |
| Jun | 4 | 72 |
| Jul | 3.33333 | 55 |
| Aug | 2 | 102.917 |
| Sep | 4 | 132.25 |
| Oct | 32.6667 | 563.5 |
| Nov | 118.667 | 631.2 |
| Dec | 75 | 409.8 |

Mean number of flies

|  | B. neohumeralis | B. tryoni |
| --- | --- | --- |
| Jan | 51.6667 | 158.8 |
| Feb | 21.3333 | 57.4 |
| Mar | 49 | 96.0667 |
| Apr | 16.3333 | 29.5 |
| May | 7.66667 | 23 |
| Jun | 15.6667 | 29.5 |
| Jul | 16.6667 | 66.5 |
| Aug | 37 | 195.5 |
| Sep | 59.3333 | 169.083 |
| Oct | 42 | 140.2 |
| Nov | 73.6667 | 312.667 |
| Dec | 109.667 | 389.267 |

Percentage of total trap catches

|  | B. tryoni | B. neohumeralis |
| --- | --- | --- |
| Jun | 2 | 1 |
| Jul | 1 | 0 |
| Aug | 3 | 1 |
| Sep | 21 | 13 |
| Oct | 18 | 11 |
| Nov | 11 | 16 |
| Dec | 4 | 17 |
| Jan | 3 | 6 |
| Feb | 6 | 14 |
| Mar | 18 | 15 |
| April | 9 | 5 |
| May | 2 | 1 |

|  | B. tryoni | B. neohumeralis |
| --- | --- | --- |
| Jun | 0 | 0 |
| Jul | 0 | 0 |
| Aug | 1 | 0 |
| Sep | 2 | 1 |
| Oct | 7 | 3 |
| Nov | 19 | 22 |
| Dec | 72 | 53 |
| Jan | 1 | 2 |
| Feb | 0 | 15 |
| Mar | 0 | 7 |
| April | 0 | 0 |
| May | 0 | 0 |

|  | Percentage of total trap catches |
| --- | --- |
| B. tryoni | 0 |

Mean number of flies (B. tryoni)

B. neohumeralis

|  | B. tryoni | B. neohumeralis |
| --- | --- | --- |
| Jan | 768.286 | 22.3333 |
| Feb | 1057.14 | 31 |
| Mar | 1257 | 22.3333 |
| Apr | 903.238 | 10.6667 |
| May | 748.095 | 0 |
| Jun | 238.857 | 0.666667 |
| Jul | 96.1429 | 1 |
| Aug | 141.19 | 1 |
| Sep | 724.286 | 10.6667 |
| Oct | 785.625 | 15.3333 |
| Nov | 1031.5 | 24.6667 |
| Dec | 953.375 | 12 |

Percentage of total trap catches

|  | B. tryoni | B. neohumeralis |
| --- | --- | --- |
| Jun | 1 | 0 |
| Jul | 1 | 0 |
| Aug | 1 | 1 |
| Sep | 26 | 26 |
| Oct | 19 | 33 |
| Nov | 6 | 6 |
| Dec | 4 | 2 |
| Jan | 31 | 31 |
| Feb | 4 | 3 |
| Mar | 4 | 1 |
| April | 6 | 1 |
| May | 4 | 0 |

Mean number of flies

|  | B. neohumeralis | B. tryoni |
| --- | --- | --- |
| Jan | 41.6667 | 165.667 |
| Feb | 40.3333 | 81.6667 |
| Mar | 27 | 133 |
| Apr | 27 | 106 |
| May | 4 | 37.3333 |
| Jun | 10.3333 | 32.3333 |
| Jul | 6.66667 | 27 |
| Aug | 23.6667 | 136.25 |
| Sep | 128 | 811 |
| Oct | 39.3333 | 219.75 |
| Nov | 24.6667 | 49 |
| Dec | 17 | 46.3333 |

Number of flies

|  | B. tryoni | B. neohumeralis |
| --- | --- | --- |
| Jan-Mar | 41 | 78 |
| Apr-Jun | 49 | 141 |
| July-Sep | 15 | 1 |
| Oct-Dec | 28 | 2 |

Percentage of total trap catches

|  | B. tryoni | B. neohumeralis |
| --- | --- | --- |
| Jun | 2 | 4 |
| Jul | 2 | 3 |
| Aug | 9 | 15 |
| Sep | 31 | 23 |
| Oct | 17 | 8 |
| Nov | 8 | 6 |
| Dec | 2 | 5 |
| Jan | 12 | 11 |
| Feb | 13 | 16 |
| Mar | 4 | 6 |
| April | 1 | 2 |
| May | 1 | 2 |

**Figure S4.** Seasonal abundance data collected by May (1961).

Number of flies

|  | B. tryoni | B. neohumeralis |
| --- | --- | --- |
| 2013-02-01 | 413 | 413 |
| 2013-03-01 | 1127 | 0 |
| 2013-05-01 | 632 | 0 |
| 2013-06-01 | 24 | 0 |
| 2014-02-01 | 277 | 61 |
| 2014-03-01 | 1259 | 764 |
| 2014-04-01 | 185 | 0 |
| 2014-05-01 | 100 | 0 |
| 2014-06-01 | 181 | 5 |
| 2014-07-01 | 89 | 0 |
| 2014-08-01 | 103 | 0 |
| 2014-09-01 | 20 | 0 |
| 2015-02-01 | 43 | 17 |
| 2015-03-01 | 3302 | 2828 |
| 2015-04-01 | 11 | 0 |
| 2015-06-01 | 37 | 0 |
| 2015-07-01 | 41 | 0 |
| 2015-08-01 | 160 | 42 |

Number of flies

|  | B. tryoni | B. neohumeralis |
| --- | --- | --- |
| 2013-03-01 | 67 | 0 |
| 2013-09-01 | 278 | 278 |
| 2014-09-01 | 33 | 15 |
| 2014-10-01 | 319 | 319 |
| 2004-11-01 | 3 | 0 |
| 2015-08-01 | 0 | 3 |
| 2015-09-01 | 116 | 110 |
| 2015-10-01 | 19 | 19 |

Number of flies

|  | B. tryoni | B. neohumeralis |
| --- | --- | --- |
| 2013-11-01 | 17 | 0 |
| 2014-08-01 | 12 | 0 |
| 2014-09-01 | 78 | 0 |
| 2014-10-01 | 218 | 157 |

Number of flies

|  | B. tryoni | B. neohumeralis |
| --- | --- | --- |
| 2013-09-01 | 505 | 0 |
| 2013-10-01 | 1189 | 0 |
| 2013-11-01 | 64 | 0 |
| 2014-09-01 | 10565 | 6716 |
| 2014-10-01 | 10365 | 7099 |
| 2014-11-01 | 1650 | 134 |
| 2014-12-01 | 327 | 0 |
| 2015-08-01 | 54 | 0 |
| 2015-09-01 | 8 | 0 |
| 2015-10-01 | 10 | 0 |

Number of flies

|  | B. tryoni | B. neohumeralis |
| --- | --- | --- |
| 2013-10-01 | 71 | 33 |
| 2013-11-01 | 28 | 0 |
| 2014-10-01 | 431 | 431 |
| 2014-11-01 | 163 | 164 |
| 2015-09-01 | 26 | 0 |
| 2015-10-01 | 7 | 7 |

Number of flies

|  | B. tryoni | B. neohumeralis |
| --- | --- | --- |
| 2012-12-01 | 491 | 0 |
| 2013-01-01 | 140 | 0 |
| 2013-11-01 | 1107 | 700 |
| 2013-12-01 | 353 | 0 |
| 2014-11-01 | 377 | 377 |
| 2014-12-01 | 2614 | 2149 |
| 2015-01-01 | 90 | 0 |
| 2015-02-01 | 75 | 0 |

**Figure S5**. Host fruit usage on fruits where both *B. tryoni* and *B. neohumeralis* emerged.
